# Evidence of distal regulations orchestrated by RNAs initiating at short tandem repeats

**DOI:** 10.1101/2025.04.10.648102

**Authors:** Mathys Grapotte, Christophe Vroland, Diego Garrido-Martín, Alessio Vignoli, Lisa Calero, Quentin Bouvier, Mathilde Robin, FANTOM consortium, Chi Wai Yip, Piero Carninci, Clément Chatelain, Laurent Bréhélin, Cédric Notredame, Roderic Guigo, Charles-Henri Lecellier

## Abstract

Short Tandem Repeats (STRs), also called microsatellites, correspond to tandemly repeated short DNA motifs (1 to 6 bp) and are one of the most polymorphic and abundant repetitive elements in the human genome. Variations of their length (i.e. number of consecutive repeats) have been implicated in gene expression regulation (termed expression(e)STRs). Using cap-trapping followed by long read sequencing, we discovered that STRs can host transcription start sites, the presence of which depends mainly on STR flanking sequences. Here, we investigate the effect of SNPs located in these sequences and ask whether STR-initiating RNAs have regulatory potential. First, we develop fully interpretable deep learning-based models, called Modular Neural Networks, able to predict, for each STR class, the level of RNAs using 101bp-long sequences encompassing STRs. Analysis of MNN filters allows us to identify multiple regulatory elements and candidate transcription factors. Second, leveraging genome sequencing and gene expression data from the Genotype-Tissue Expression project, we use the output of our models to link the levels of STR-initiating RNAs to the expression of nearby genes. We identify 14,340 significant associations (coined RNA(r)STRs) and illustrate how this novel resource can help interpret non-coding variants associated with complex traits and diseases. Third, we unveil an intricate transcriptional interplay between STR-initiating RNAs and Alu repeats that may couple their regulatory actions, extending both the nature and the functional importance of non-coding transcription and shedding new light on the complexity of distal regulations orchestrated by repeated sequences.

## Introduction

Almost half of the human genome is scattered with repetitive sequences that are tandemly arrayed or interspersed ^1, 2^. More than 70% of all nucleotides are thought to be transcribed at some point ^3–5^. As a consequence, a large portion of non-coding RNAs derives from repetitive elements ^2, 6–8^ and microsatellites, also called Short Tandem Repeats (STRs), are no exception. STRs correspond to tandemly repeated DNA motifs of 1 to 6 bp (which distinguish STR class) and constitute one of the most polymorphic and abundant repetitive elements (*→* 5% of the human genome) ^9^. STRs are known to widely impact gene expression and to contribute to expression variation ^10–12^. At the molecular level, STRs can for instance affect expression by inducing inhibitory DNA structures ^13^ and/or by modulating TF binding ^9, 14–16^. Specific expression Quantitative Trait Loci (eQTLs) ^11, 12^, called expression(e)STRs ^11^, have been calculated to correlate STR length variations with the expression of nearby genes. The Cap Analysis of Gene Expression (CAGE) technology revealed that transcription start sites (TSSs) are present in diverse repetitive sequences ^7, 8^. Combining cap trapping and long-read MinION sequencing (CFC-seq) ^17^, we specifically observed that thousands of STRs, from different classes, can initiate transcription in a variety of human and mouse cells and produce a variety of transcripts (including protein-coding, long non-coding and enhancer RNAs). Consistent with our results, the oncogenic transcription factor EWS-FLI1 triggers bidirectional transcription at (*GGAA*)*_n_*STRs in Ewing sarcoma cells ^18^. We also observed that STRs associated with clinically relevant variants correspond to those associated with high CAGE signal, suggesting functional role(s) for this STR-associated transcription ^17^

Here, we develop a method to evaluate, at a large scale, the regulatory potential and the functional consequences of STR-initiating RNAs. STRs are often if not invariably studied in population genetics through variations of their lengths ^11, 12, 19^. However, we previously revealed key roles of STR flanking sequences to predict transcription initiation and RNA level ^17^. In addition, we observed that genetic variants linked to human diseases, are located around STRs associated with high RNA levels. Hence, when considering the potential regulatory impact of STR-initiating RNAs, STR length may not be the sole relevant feature, as previously suspected ^20^. In fact, in Ewing sarcoma cells, the number of (*GGAA*)*_n_* demonstrates no length-dependency with EWS-FLI responsiveness for gene repression, as opposed to gene activation ^21^. Considering STRs and SNPs located at their vicinity, genetic elements which have been called SNPSTRs, provides complementary evolutionary information and improves genetic studies ^22, 23^. SNPs located within a 50kb window of the STR can also be used to infer STR genotypes ^24^. At the molecular level, SNPs located in STR surrounding sequences can act in complex relationships with STR length variation to control gene regulation ^25^. Together with ours, these findings suggest that STR-flanking sequences and STR-surrounding SNPs should be considered to fully appreciate the impact of STRs on gene expression.

To predict the level of RNAs initiating at STRs, we initially developed Convolutional Neural Networks (CNNs) ^17^. Although methods exist to interpret this type of models after learning ^26–28^, CNN interpretation still has some limitations ^29–32^, making it hard to evaluate the biological significance of their predictions. To circumvent these limitations but still be able to consider the impact of STR-surrounding SNPs, we develop fully interpretable modular neural networks (MNNs), which combines learning and interpretation in one single step preserving automatic feature extraction from DNA sequence encompassing STRs. Contrary to other methods ^26–28^, the interpretation step of MNNs is integrated in the learning procedure and takes the whole set of sequences, not a subset of specific inputs. Analyses of MNN filters allow us to identify several regulatory elements and transcription factors governing the expression of STR-initiating RNAs. We further paired STRs with nearby genes and use MNN predictions to regress gene expression, thereby computing new type of expression(e)STRs, coined RNA(r)STRs (to refer to the regulatory potential of RNAs initiating at STRs), which links the level of RNA initiating at STRs and the expression of nearby genes. Our rSTR catalog is enriched in previously identified eQTLs ^33^ and eSTRs computed solely on STR length variations ^34^ but also provide novel STR-gene regulatory associations. We also unveil a complex transcriptional interplay between rSTRs and Alu elements. Together, our results provide evidence that STR-mediated gene regulation can also rely on RNA molecules acting distantly, thereby extending both the nature and the functional importance of non-coding transcription ^35–38^. We make our rSTR catalog publicly available so that it can serve as a useful resource for the interpretation of genetic variants located at the vicinity of STRs.

## Results

### Predicting RNA levels with Modular Neural Networks

The modular neural network (MNN) is a fully explainable deep neural network, which identifies DNA motifs correlated with RNA levels (Figure 1A). It takes as input 101-bp long sequences centered around STR 3’ end as defined by the strand of the transcription (roughly the size of the core promoter ^39^). The predicted signal corresponds to the CAGE signal averaged across all samples and normalized by the length of the STRs as in ^17^. The normalization by STR length was motivated not only by the impact of very low CAGE counts along STRs, which artificially increase the signal, but also by the results of GTEx long reads sequencing ^40^, revealing that the levels of transcripts initiating at STRs are not correlated with STR length (Supplementary Figure S1). Moreover, the Gini coefficient of STR-initiating transcripts is lower than that observed for all transcripts (Supplementary Figure S2), suggesting that this type of RNAs is expressed more ubiquitously than others. Hence, MNNs focused on cell-agnostic and initiation-specific events. The network architecture is determined dynamically during training by adding modules one after the other. Each module encapsulates a single convolution filter layer followed by a ReLU activation and a regression layer. ReLU will output the input directly if it is positive, or output zero otherwise. One module is characterized by its convolution filter size. Finally, a global regression layer takes as input the output of each module and yields a final prediction. Each module learns a motif information encapsulated in the convolution filter and a position information encapsulated in the module regression layer. The simplicity of the design ensures that the model is fully explainable. The learned convolution filter can be assimilated to a Position Weight Matrix (PWM) ^41^ and the module regression layer weights to the PWM expected location. The MNN is trained dynamically and asynchronously using a custom training algorithm: each module harbors different hyperparameters (convolution filter size, momentum and learning rate) and modules are added and trained one-by-one until no further improvement is measured using an early stopping strategy. Module weights are frozen once the training of this particular module has stopped. Such strategy prevents the model from re-adapting module weights and forces new modules to learn different motifs/position combination to ensure performance improvement and limit motif overlap. To leverage the fewer amount of MNN parameters (which makes it hard to learn complex distribution) and in order to improve generalization, we linearized the CAGE signal and predicted ranks instead of CAGE signal ^17^. Thereby, the distribution of observed values become consistent across STR classes: the ranked signal being transformed in an affine function of slope 1, it follows exactly the rectified linear unit (ReLU) activation function for x *>* 0, which improves precision and convergence speed. MNNs yielded overall good performances with Spearman correlation coefficient between observed and predicted ranks *>* 0.5 for several classes (Figure 1B and Supplementary Figure S3). MNNs used between 4 (for (*AG*)*_n_* STR class) and 10 (for (*AAT*) *_n_* STR class) filters (Figure 1C), showing that few filters are needed to reach the observed performances. We also learned MNN models using as input 101-bp long sequences aligned on STR 5’end (as defined by the strand of the transcription) but these models were found inaccurate (Spearman rho *→* 0).

**Figure 1.**
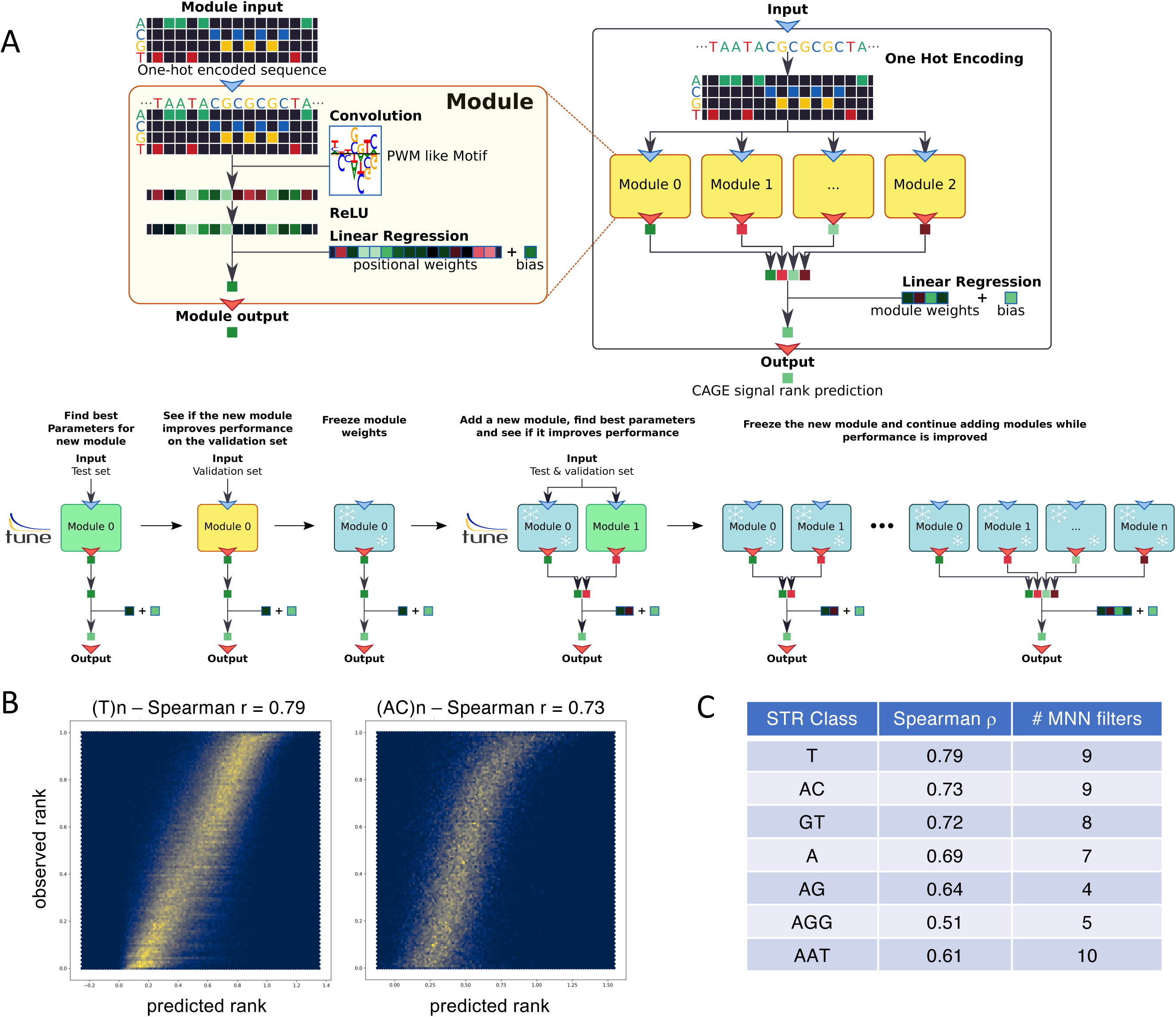
Predicting CAGE signal at STRs using Modular Neural Networks (MNNs).a. Principles of Modular Neural Networks. see text for details **b.** Hexbin plots showing predicted (y-axis) vs. observed ranks (x-axis) for (*T*) *_n_* and (*AC*)*_n_* STRs. The Spearman *ε* correlation is indicated. **c.** MNN performances measured as Spearman *ε* correlation between observed and predicted ranks for different STR classes. The numbers of filters learned by MNNs are indicated. Note that the CNN models initially built have 110 filters 17.

### Identification of candidate transcriptional regulators of STR-initiating RNAs

To better understand the molecular basis governing the expression of STR-initiating RNAs, we compared the filters identified by MNNs to known TF motifs. Note that we considered the sequences of 22,491 STRs corresponding to STR with prediction error on the reference genome ↭ 0.05. For each 101bp-long sequence centered around STR 3’end (strand being defined as strand of transcription), we retrieved foreground and background subsequences (ReLU *>* 0 and =0 respectively) and ran HOMER for each module of each STR-specific MNN model. HOMER motifs were compared to known JASPAR PWMs (JASPAR2022 core vertebrate ^42^ and JASPAR2020 POLII ^43^ collections) identifying several TFs potentially regulating transcription at STRs (detailed results are available at https://gite.lirmm.fr/ml4reggen/rstr). Restricting to matches *>* 40% in the foreground sequences and HOMER identity score *>* 0.7 (Supplementary Table S1A), we found that the C2H2-type zinc finger ZNF384 was the top TF present in 11 out of 43 STR models, followed by 3 downstream core elements (DCEs: POL008.1 present in 8, models POL0010.1 and POL009.1 present in 7 models) and the initiator motif (INR, POL002.1 present in 6 models) (Supplementary Table S1A and Figure 2), all members of the core promoter elements implicated in transcription initiation ^39^. Several short (typically *<* 3bp) MNN filters did not match a PWM (Supplementary Figure S4), in agreement with ^44, 45^. The characterization of three DCEs is in agreement with findings supporting a key role of downstream promoter region in human gene transcriptional regulation ^46, 47^. However, because (i) CAGE measures RNA abundance and (ii) the region downstream TSS also corresponds to RNA sequence, we could not exclude that these motifs are also implicated in post-transcriptional regulations. The STRs considered here are dominated by A/T-rich STRs (Supplementary Figure S5) and the 10 models wherein ZNF384, whose PWM corresponds to a poly(A)/poly(T) stretch (MA1125.1), appears important (namely (*A*)*_n_*, (*AAAAAC*)*_n_*, (*AAAC*)*_n_*, (*AAT*) *_n_*, (*AATT*) *_n_*, (*ATTT*) *_n_*, (*CTTT*) *_n_*, (*CTTTT*) *_n_*, (*GTTTT*) *_n_* and (*T*) *_n_*) cover 64% (14,470) of all 22,491 STRs tested. As shown Figure 2B, the activation of the filter corresponding to ZNF384 coincides with the end of the STRs. To assess the specificity of each filter of each STR class model, we designed a second strategy: for a given foreground corresponding to one filter of one model, all sub-sequences with ReLU *>* 0 obtained for all other filters of the same model were used as background. In that case, limiting the output to matches *>* 40% in the foreground sequences and HOMER identity score *>* 0.7, ZNF384 remains the top TF present in 18 out of 43 models (Supplementary Table S1B) and, again, these models correspond to A/T-rich STRs (namely (*A*)*_n_*, (*AAAAAT*) *_n_*, (*AAAAC*)*_n_*, (*AAAAG*)*_n_*, (*AAAC*)*_n_*, (*AAAG*)*_n_*, (*AAAT*) *_n_*, (*AAT*) *_n_*, (*ATT*) *_n_*, (*ATTC*)*_n_*, (*ATTT*) *_n_*, (*ATTTT*) *_n_*, (*CT*) *_n_*, (*CTTT*) *_n_*, (*CTTTT*) *_n_*, (*GTT*) *_n_*, (*GTTTT*) *_n_*, (*T*) *_n_*). We further confirmed the preferential binding of ZNF384 at A/T-rich STRs by leveraging ZNF384 ChIP-seq data collected in various cell types (GM12878, HEK293T, Hep-G2 and K-562) using the Remap database ^48^ (Figure 2C). Similar results were obtained by the Codebook consortium ^49^. We also evaluated the functional consequences of ZNF384 knock-down (KD) on STR transcription using ENCODE CRISPRi experiment (ENCSR103URL and ENCSR932JIP). The read count was made on annotated transcripts (ENCODE and FANTOM) as well as RNAs detected by Cap-trap full-length cDNA sequencing (CFC-seq) ^50^ (Supplementary Table S2). 9,987 out of 41,766 genes with at least one transcript initiating in a window of 5bp around an STR were significantly modulated (adjusted p-value *<* 0.05 in Supplementary Table S2), while, it is true for 2,087 out of 11,800 genes without transcript initiating around STRs (odds ratio 1.5, Fisher’s exact test p-value *<* 2.2e-16). The median fold change for STR-initiating genes was negative (−2.5), indicating an overall positive action of ZNF384 on STR-initiating transcription.

**Figure 2.**
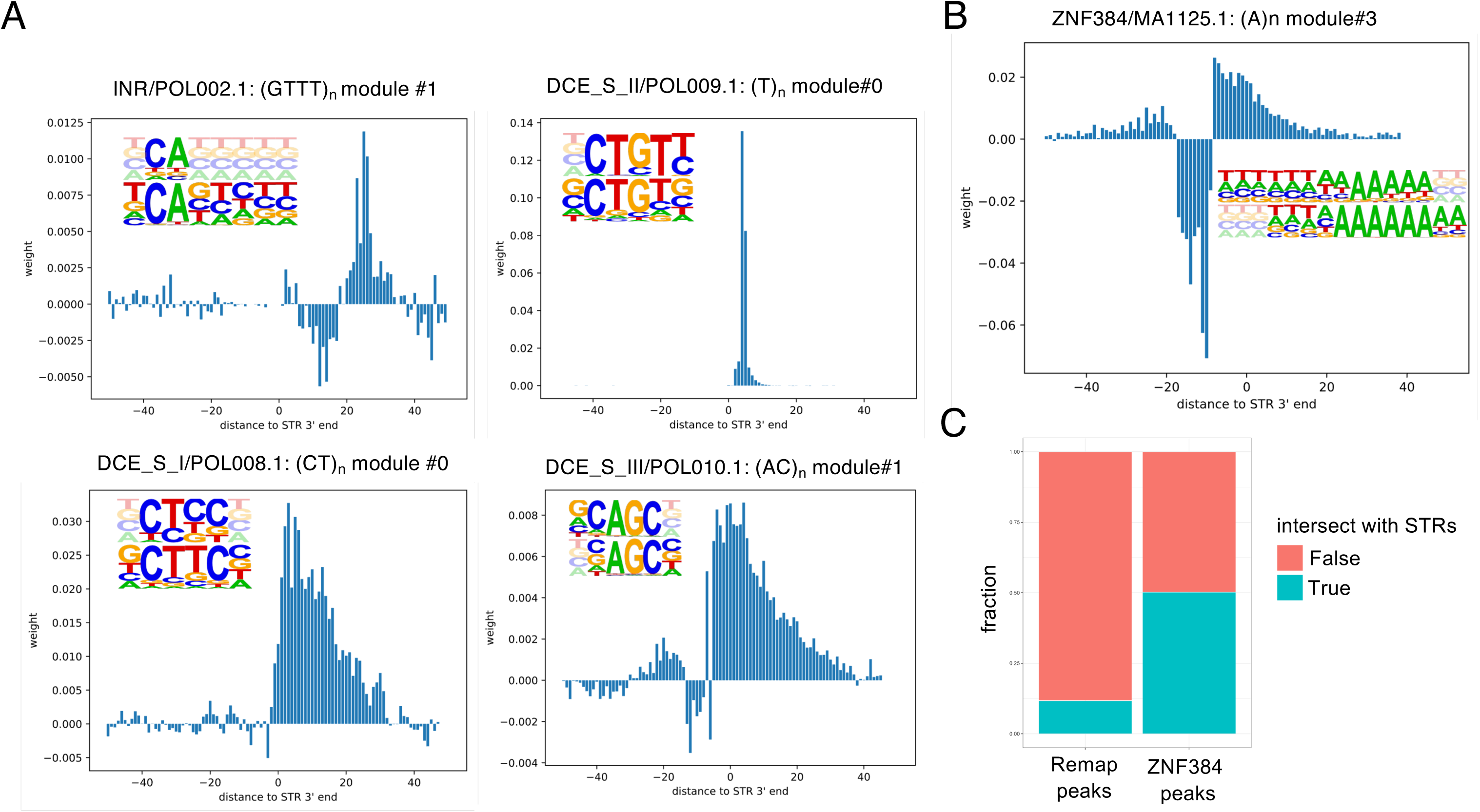
MNN filter analyses. **a.** Examples of filters corresponding to promoter core elements. The distribution of filter activations around STR 3’end is shown. Activation corresponds to the multiplication of the mean of the activation score after the ReLU layer for all sequences of the model by the weights of the first regression layer and by the weight of the module in the second regression layer. Alignment between HOMER (top) and JASPAR (bottom) motifs is also indicated. For each filter, the motif name, JASPAR ID, STR class and module number are indicated. Detailed results are provided in Supplementary Table S1A and at https://gite.lirmm.fr/ml4reggen/rstr **b.** Example of filter corresponding to ZNF384 motif. Results are presented as in a. **c.** Fraction of Remap ChIP-seq peaks (left) or ZNF384 ChIP-seq peaks (right) intersecting (blue) or not (red) with STRs. 42,049 ZNF384 peaks out of 83,673 (*→* 50%) intersect with STRs, while it is true for 8,063,480 out of all 68,655,741 Remap TF peaks (*→* 12%, odds ratio *→* 7.59, Fisher’s exact test p-value *<* 2.2e-16).

### A novel RNA(r)STR catalog

We further investigated whether STR-initiating RNAs have a regulatory potential by assessing their impact on gene expression. Specifically, we regressed gene expression on the output of our MNN models, which predict the level of STR-initiating RNAs. These regressions, coined rSTRs, assess the potential relationship between the level of transcription at STRs and the expression of nearby genes. Note that RNA-seq depth is not sufficient to properly quantify STR-initiating RNA levels and rSTR computations necessitate MNN predictions. For sake of comparison with previous eSTR computations, we considered GTEx V7 data and paired genes with STRs located in a 100kb window of 100kb ^34^. Predictions were computed for each STR and for each individual, inserting SNPs of each allele located in a window of 50bp around the STR 3’end (as defined by the transcription strand) in the input sequence (several SNPs can be considered for each individual sequence). Only STRs with prediction error on the reference genome, measured as (predicted rank - observed rank) ↭ 0.05 and variance of MNN scores in GTEx data ↫ 10*^→^*3, were considered. Each STR was then associated with two prediction scores (one for each allele) for each individual. The two scores (as a multivariate response) were jointly regressed on gene expression and other co-variates such as gender, age and two first principal components (PCs) resulting from principal components analysis (PCA) on SNP genotypes from each sample using the MANTA method ^51^. The whole pipeline (Figure 3), from SNP insertion into sequences to MNN prediction and rSTR computations, was made available as a Nextflow workflow ^52^ (https://gite.lirmm.fr/ml4reggen/rstr). Because we used thresholds on model errors and variance, the STR catalog tested here is limited to certain STR classes (Supplementary Table S3). A total of 115,608 (STR, gene) pairs were tested and MANTA yielded 14,340 significant associations between 8,458 rSTRs and 11,060 target genes (rGenes) in at least one tissue at a false discovery rate (FDR) of 5%: 78% of genes are associated with only 1 STR and 60% of STRs with only 1 gene i.e. median of 1 STR associated with 1 gene and 1 gene associated with 1 STR (Supplementary Tables S4A and S4B).

**Figure 3.**
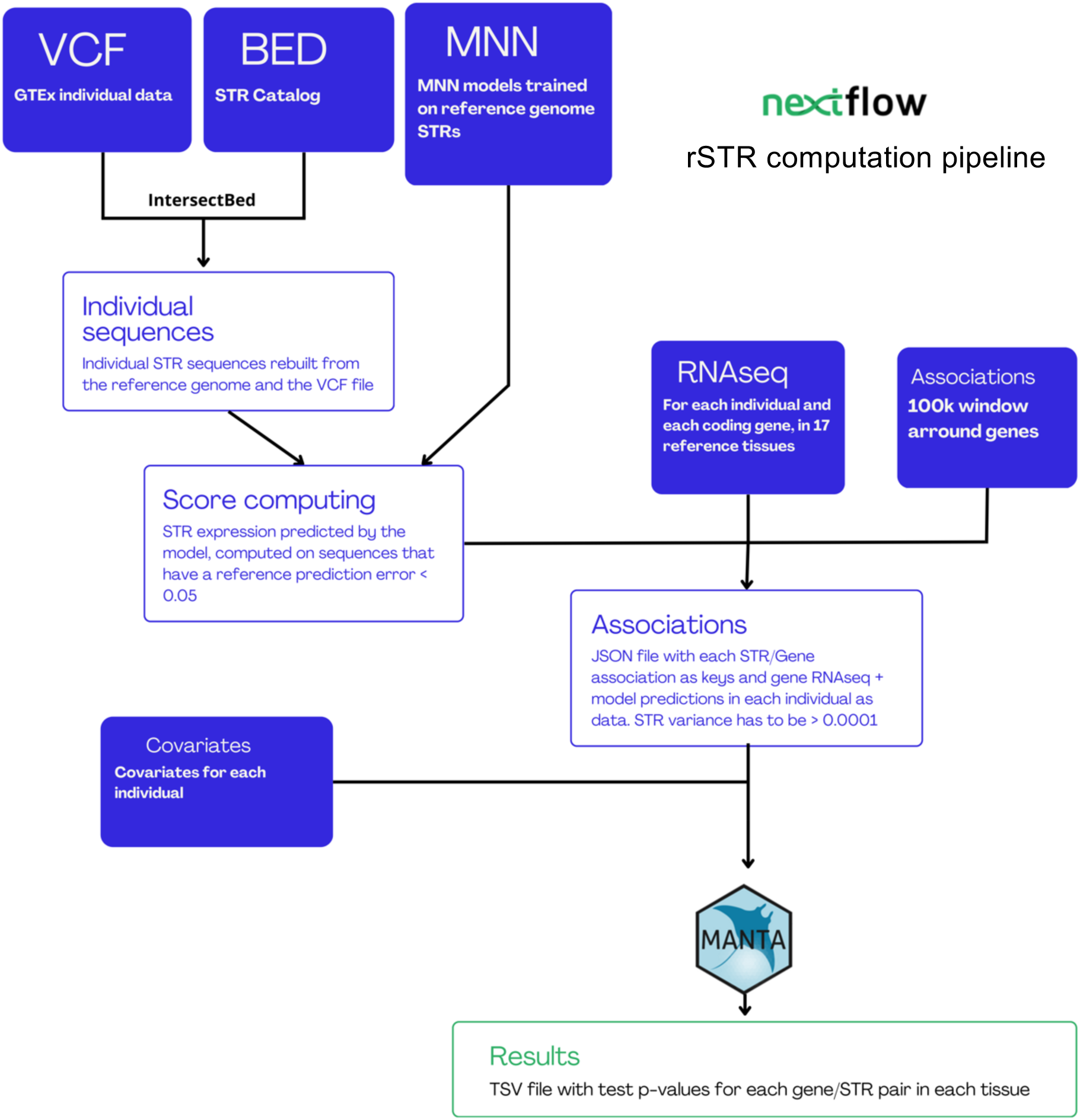
rSTR computation pipeline. Required input files are indicated as blue boxes. The SNPs located in a window of 50bp around STR 3’end (STR strand being defined by CAGE signal - ‘STR catalog’ blue box) were retrieved from GTEx V7 individual data (’VCF’ blue box). These SNPs were further introduced into the corresponding STR reference sequence and MNN prediction scores were generated using a custom-made python script available at https://gite.lirmm.fr/ml4reggen/rstr (’MNN’ blue box). Each STR was associated with two prediction scores (one for each allele) for each individual. Genes and STRs were paired considering each STR located in a window of 100kb around GENCODE V19 genes (’Associations’ blue box). The two scores of each STR (considered as a multivariate response) were jointly regressed on GTEx V7 gene expression (RNAseq blue box) and other co-variates such as gender, age and two first principal components (PCs) resulting from principal components analysis (PCA) on SNP genotypes from each sample (’Covariate’ blue box) using the MANTA method 51. The whole pipeline was made available as a Nextflow workflow 52 (https://gite.lirmm.fr/ml4reggen/rstr)

### Evaluation of rSTR significance

A strong enrichment in eQTL associations was noticed among the rSTR pairs identified: 1,550 (STR, gene) pairs out of the 115,608 tested (*→* 1.3%) and 556 rSTRs out of the significant 14,340 (*→* 3.8%) were also identified as GTEx eQTLs ^33^ (hypergeometric test p-value *<* 2.2e-16). Likewise, 595 pairs out of the 115,608 tested (*→* 0.5%) and 208 rSTRs out of the significant 14,340 (*→* 1.4%) were previously reported as eSTRs computed solely with STR length variation ^34^ (hypergeometric test p-value *<* 2.2e-16), in line with the observations that, compared to all STRs, those closest to transcription start sites and near DNase I hypersensitive (HS) sites are more likely to be robust fine-mapped eSTRs ^34^. Thus, rSTRs computed with MNN predictions scores recover a fraction of eQTLs ^33^ as well as eSTRs computed by length variation ^34^, but also identify new associations undetected by both approaches. We also noticed that 26 rSTRs out of 14,340 (*→* 0.2%) and 56 (STR, gene) pairs out of the 115,608 tested (*→* 0.05%) correspond to splicing QTLs ^53^ (hypergeometric test p-value*→* 6e-11), indicating that some rSTRs could exert their regulatory action at the post-transcriptional level. Note that, among these 26 rSTRs, only 5 also correspond to eQTLs. Despite these enrichments, the vast majority of the 14,340 rSTRs detected correspond to novel associations (n = 13,190, *→* 92%). To confirm the existence of physical interactions between STR-initiating RNAs and distal genes, we first looked at rSTR associations found in DNA-DNA interactions mediated by RNA polymerase II (Pol II) and detected by ChIAPET. Specifically, the fraction of significant rSTRs (Supplementary Table S4) detected among all possible (rSTR, rGene) pairs formed by ChIA-PET was compared to the fraction of all tested rSTRs (Supplementary Table S3) obtained with all possible (tested STR, tested gene) pairs using Fisher’s exact tests in 5 different cell types (see Methods section). We invariably observed a significant enrichment of rSTRs in all ChIA-PET datasets (Figure 4A). Second, we reasoned that some interactions between STR-initiating RNAs and target genes should be detectable by RNA-chromatin interaction data. We leveraged RNA-DNA interaction RADICL-seq data collected in 19 different conditions by the FANTOM consortium (Lambolez*et al.*, in preparation). The RNA part of RADICL-seq reads were mapped to known transcripts (ENCODE or FANTOM) and/or RNAs detected by cap-trapped long read ONT RNA sequencing (ONT-CAGE / CFC-seq) ^50^ whose 5’ ends were located in a window of 5bp around STRs. The DNA part of RADICL-seq reads were intersected with gene coordinates. For comparison, we defined a set of non-rSTRs as, among the tested pairs, those implicating STRs never found associated with the expression of any gene and genes whose expression was never found associated with STR transcription (n = 36,199). Hence, 63 rSTRs and 172 non-rSTRs could be compared. Among these pairs, 29 out of 63 and 41 out of 172 were detected by RADICL-seq, revealing an enrichment of rSTRs compared to non-rSTRs in RNA-DNA interactions (odds ratio = 2.71, Fisher’s exact test p-value = 1e-3). Note that comparing fraction of rSTRs detected in RADICL-seq to that obtained with all tested rSTRs yielded non significant results, likely due to the presence of potential false negative rSTRs ^54^. Interestingly, 22 of these 29 rSTRs (*→* 76%) were detected in 2 or more RADICL-seq libraries (Figure 4B). Note that, at this stage, it is not possible to evaluate the cell-specificity of rSTRs simply because MNN predictions are cell-agnostic (the same initiation level is predicted for a given STR for all tissues of one individual).

**Figure 4.**
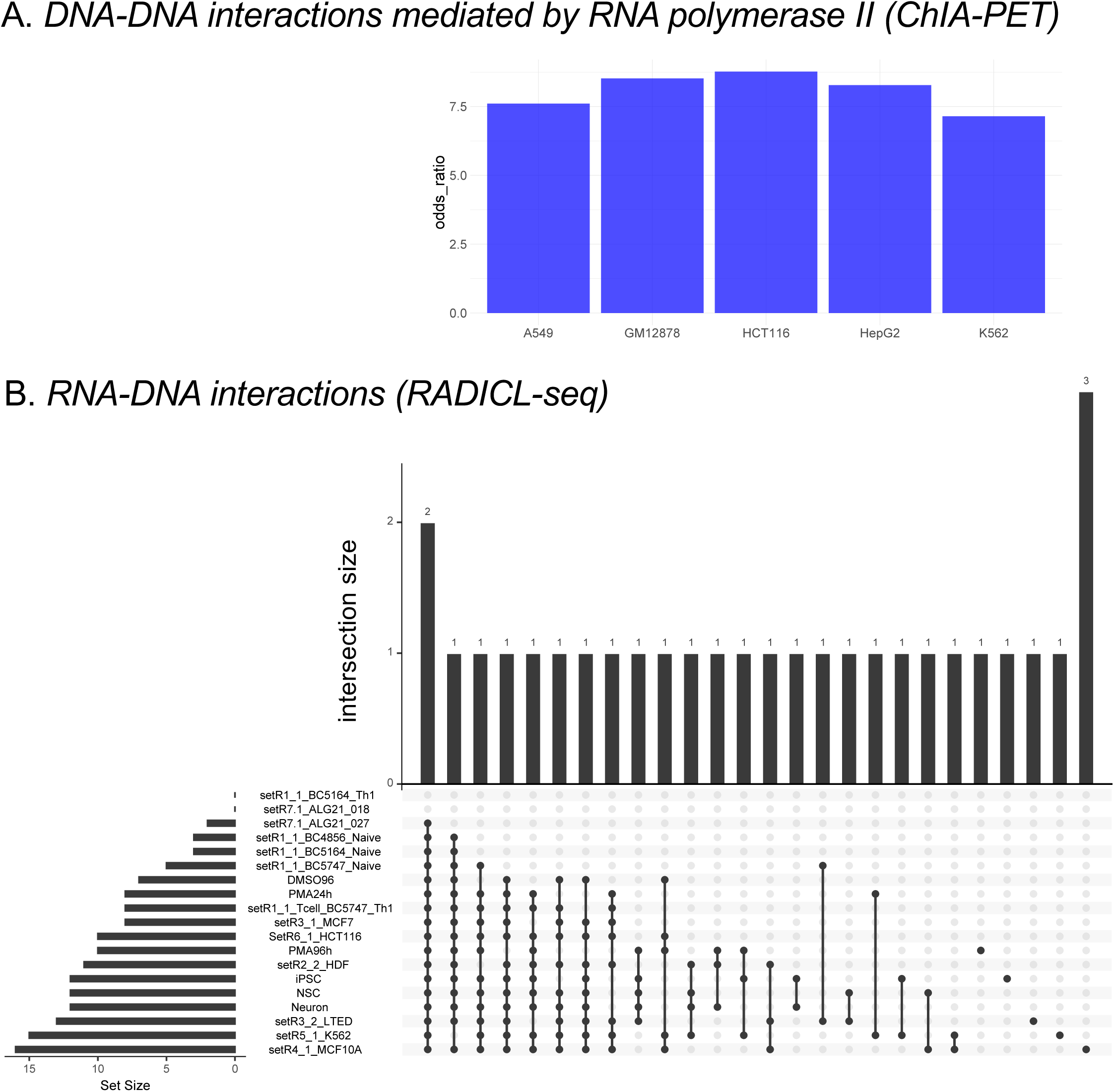
Evaluating rSTR biological relevance. **a.** The fraction of rSTRs detected among all possible (rSTR, rGene) pairs formed by ChIA-PET was compared to that obtained with all possible (tested STR, tested gene) pairs using Fisher’s exact tests in 5 different cell types. All p-values are *<* 2.2e-16. Odds ratio are shown as barplots. For each cell type, the following fractions correspond to number of rSTRs among all possible (rSTR, rGene) pairs formed by ChIA-PET and number of tested STRs among all possible (tested STR, tested gene) pairs formed by ChIA-PET. K562: 996/7015 vs. 7447/328992 ; HepG2 73/536 vs. 554/29645 ; HCT116: 239/1563 vs. 1737/86106 ; A549: 212/1554 vs. 1641/80644 ; GM12878: 264/1837 vs. 2327/120421. **b.** Distribution of the 29 rSTRs detected by RADICL-seq in 19 libraries. 22 out of 29 rSTRs (*→* 76%) are detected in 2 or more libraries.

### Interpretation of genetic variations with rSTRs

Our rSTR catalog was enriched in credible causal variants as defined in CAUSALdb ^55^: 1,728 STRs out the 8,458 engaged in rSTRs (*→* 20.4%) harbor a causal variant in their vicinity, while the same was true for 3,912 STRs among the 22,491 STRs tested (*→* 17.4%, odds ratio 1.22, one-sided Fisher’s exact test p-value= 5.4e-10). We noticed an imbalance in the distribution of causal variants around STRs engaged in rSTRs with more variants located 100bp downstream STRs than 100bp upstream (Figure 5A). The sense of STRs is given by the CAGE signal strand (Figure 5A, right). This imbalance indicated that both sides of STR are not symmetrical and could be explained, for instance, by the presence of RNA being initiated at STR, and through which causal variants can have phenotypical consequences. A similar distribution is observed when considering all tested STRs (Supplementary Figure S6), suggesting that more rSTRs have yet to be detected, as suspected above. We annotated 2,419 causal variants located 100bp downstream rSTRs and 1,177 located 100bp upstream using the scoring framework RADAR, which identifies variants potentially affecting post-transcriptional regulations and RNA-binding protein (RBP) functions ^56^. Although most causal variants scored at 0, RADAR pinpointed more variants associated with RBP-function dysregulation downstream than upstream STRs engaged in rSTRs (respectively 22 variants with score *>* 0 out of 2,419 and 3 out of 1,177, odds ratio *→* 3.59, one-sided Fisher’s exact test p-value *→* 1.7e-2, Supplementary Tables S5A and S5B). We then used CFC-seq ^50^ to identify RNAs initiating at STRs engaged in rSTRs (n=60) and RNAs initiating at STRs never engaged in rSTRs (non-rSTRs, n=96). No significant differences could be observed in terms of transcript length and CAGE signal (Figure 5B and 5C). The coordinates of these two classes of RNAs (full length, including introns) were intersected with the peak coordinates of CLIP (cross-linking and immunoprecipitation) data targeting several RBPs ^57^. We found that rSTR-initiating RNAs contain more CLIP peaks than RNAs initiating at non-rSTRs (respectively 16,301 and 2,329 ; *→* 272 peaks per rSTR-initiating RNA vs. *→* 24 peaks per RNA initiating at non-rSTRs, Figure 5D). The preferential binding of RBPs on rSTR-initiating RNAs reinforced their functional importance. The list of RBPs preferentially bound to rSTR-initiating RNAs is provided in Supplementary Table S6.

**Figure 5.**
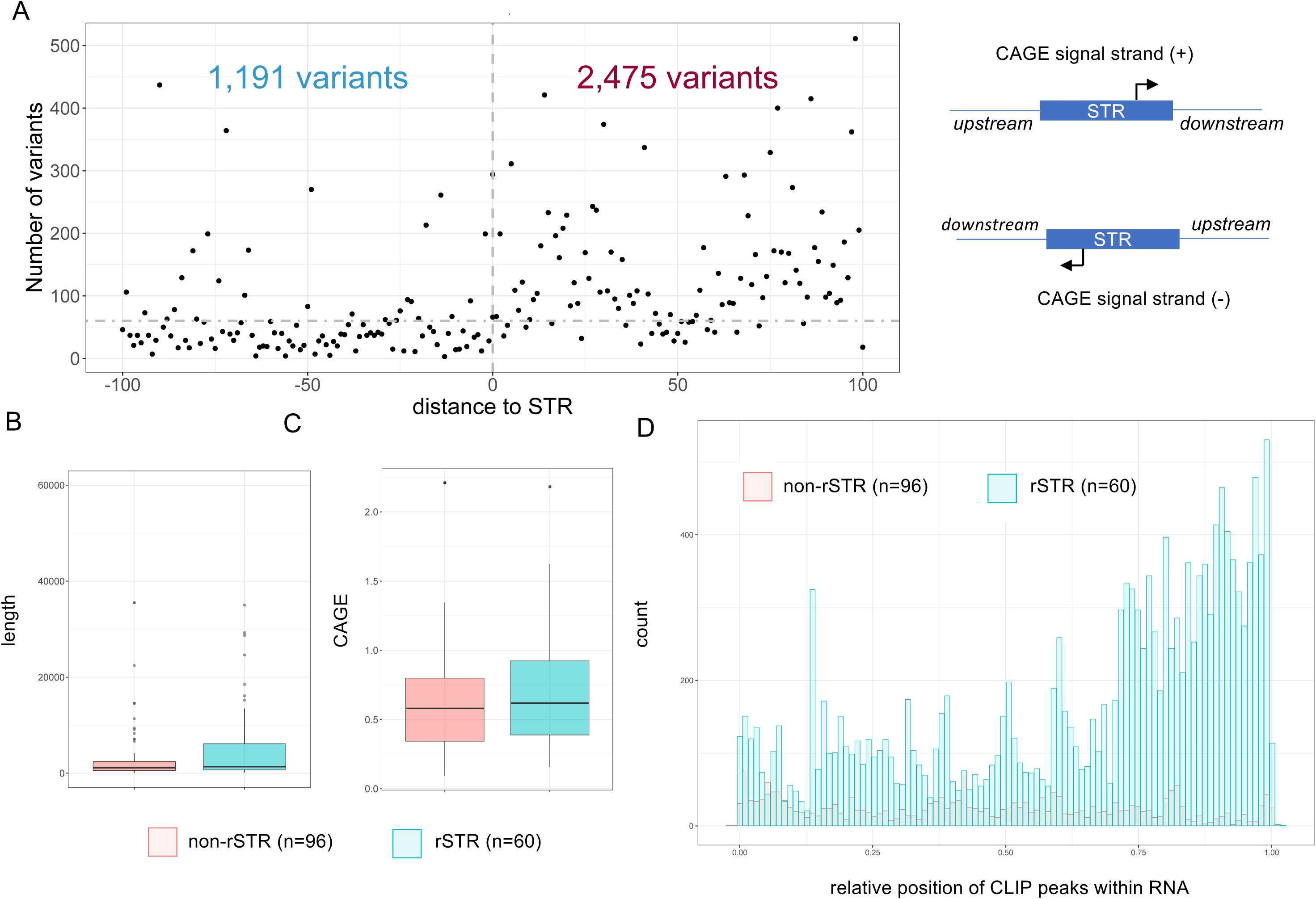
Interpreting genetic variants with rSTRs. **a.** Distribution of causal variants around rSTRs. STR sense is defined by the strand of the CAGE signal (right). Hence, one single STR associated with CAGE signal on both strand will have 2 downstream and 2 upstream sequences. x-axis: distance to STR 5’ or 3’ ends ; y-axis: number of variants (each dot is a variant). Vertical dashed grey line indicates the position of the STR. Horizontal dashed grey line is arbitrary drawn to highlight the difference between the number of variants located upstream and downstream STRs. number of variants located downstream = 2,475 ; number of variants located upstream = 1,191 ; Fisher’s exact test p-value *<* 2.2e-16, odds ratio *→* 2.53 **b.** Boxplots of length of CFC-seq-defined transcripts initiating at rSTRs (red) and non-rSTRs (blue). The lower and upper hinges correspond to the first and third quartiles (the 25th and 75th percentiles). The upper whisker extends from the hinge to the largest value no further than 1.5!IQR from the hinge (where IQR is the interquartile range or distance between the first and third quartiles). The lower whisker extends from the hinge to the smallest value at most 1.5!IQR of the hinge. Data beyond the end of the whiskers are plotted individually. Wilcoxon test p-value = 0.08755 **c.** Boxplots of CAGE signal detected at STRs engaged in rSTR (blue) and non-rSTR (red) associations. Wilcoxon test p-value = 0.2538 **d.** Relative distribution of CLIP peaks along STR-initiating RNAs. The coordinates CFC-seq-defined transcripts (including introns) were intersected with RBP CLIP peaks from POSTARDB. The distribution of relative peak positions along transcripts initiating at rSTRs (blue) and non-rSTRs (red) is shown.

### A case study of genetic variant interpretation with rSTR

We further illustrated the relevance of using rSTRs to interpret genetic variants with rs12151021, which is located in ABCA7 intron 18, 50bp downstream the Human STR 654307;AAT;+ STR (Supplementary Table S4A and S4B). Our rSTR catalog indicated that the expression of the RNA initiating at this STR is associated with that of GRIN3B (ENSG00000116032, located *→* 41kb upstream the STR) in ‘Thyroid’ tissue (Supplementary Figure S7). This gene was also found associated with rs12151021 in five tissues in GTEx V8 (https://gtexportal.org/home/snp/rs12151021), not in the V7 version used here for comparison, and this association was independently confirmed in brain cortex ^58^. Of note, GRIN3B was associated with another STR in ^34^ (Human STR 654285;AAAAT located at chr19:1025181-1025235 on hg38). At the phenotypic level, rs12151021 is linked to the ‘red cell distribution width’ and ‘Mean corpuscular hemoglobin concentration’ traits as well as Alzheimer’s disease according to causalDB and ^58^. On the other hand, GRIN3B encodes a subunit of an N-methyl-D-aspartate (NMDA) receptor, which controls synaptic plasticity as well as learning and memory functions ^59^. In mouse, GRIN3B knockout (KO) leads to increased red blood cell distribution width ^60^. The overall concordant phenotypes associated with rs12151021 on one hand and GRIN3B expression on the other hand supported a regulatory relationship. At the molecular level, rs12151021 has a moderate but significant impact on MNN predictions (see x-axis in Supplementary Figure S7). At the RNA level, rs12151021 appears to impact post-transcriptional regulations as indicated by its RADAR score (1.07), which is more consistent with the mean scores of HGMD and 1000 Genomes variants within RNA-binding proteins (respectively 1.871 and 1.337) than all variants (respectively 0.589 and 0.025) ^56^. Experimental investigations must be conducted to determine the exact mechanism of action of the RNA initiating at Human STR 654307;AAT on GRIN3B expression. Nevertheless, this example illustrates how our rSTR catalog can be used to interpret genetic variations around STRs, including from a post-transcriptional perspective ^56^.

### rSTRs genomic features

No particular STR class was found to be enriched in rSTRs (Supplementary Table S7), suggesting that, among all classes tested, virtually all could have regulatory functions. The length of STRs engaged in rSTRs was not different from that of all tested STRs (median = 18 in both cases, Wilcoxon test p-value = 0.6347), in contrast to what was observed for eSTRs computed on length variations^34^. Only 3,322 rSTRs targeted their closest gene (out of 15,294 STR/gene possible pairs, *→* 21%) and the median distance between STR and gene in rSTRs was *→* 35.4kb (Figure 6A). Overall, predictions of transcription initiation at STRs (MNN scores) were either positively or negatively correlated to rGene expression (Supplementary Tables S8A, S8B and S9), suggesting that rSTRs participate in either inhibition or activation of rGene expression. The strand of the transcribed STR did not necessarily correspond to that of the rGene (i.e. transcribed STR and target gene have the same strand in 50% of the detected associations, Supplementary Table S4B). Together these results suggested that rSTRs exert their functions mostly distantly and in a *trans* mechanism. The target genes were enriched in non-coding RNAs (microRNAs and long intergenic RNAs), while depleted in protein coding genes (Supplementary Table S10). Finally, we looked at potential link between rSTRs and known regulatory elements. The STRs engaged in rSTRs were not enriched in specific ENCODE candidate *cis*-regulatory elements (Supplementary Table S11), except promoter-like signature (PLS), reflecting that transcription initiation at STRs is associated with typical biochemical signatures ^17^.

**Figure 6.**
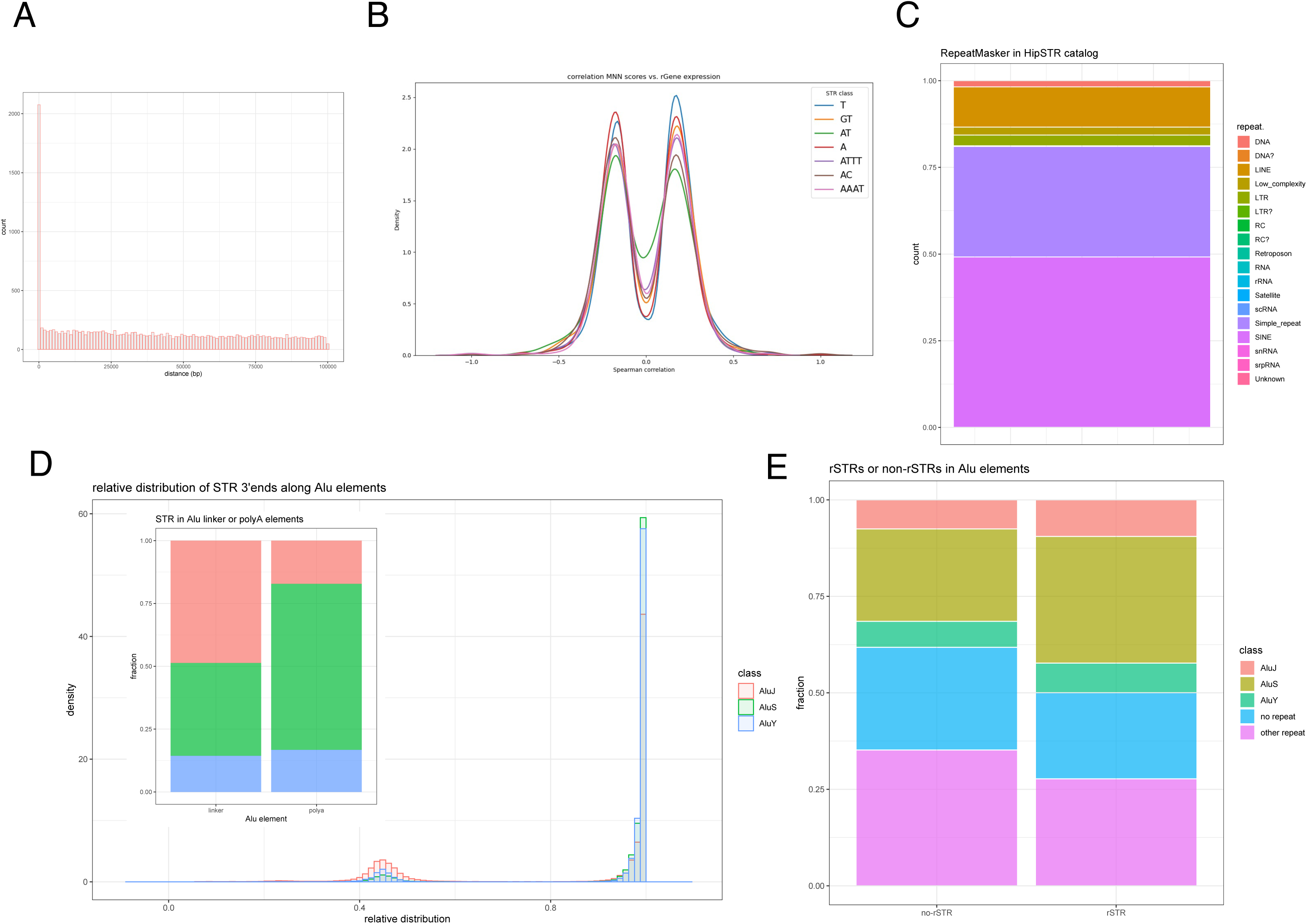
rSTR genomic features. **a.** Histogram of the distances between rSTRs and target genes (rGenes). The GENCODE V19 gene coordinates were used for computations. **b.** Distribution of Spearman correlation between MNN scores and gene expression in GTEx V7 data. For MNN scores, the average correlation obtained for both STR alleles was considered. For sake of clarity, only STR classes with more than 300 rSTRs are shown. Detailed values are provided in Supplementary Table S9. **c.** Distribution of repeat elements as defined by RepeatMasker in HipSTR STR catalog. STRs are enriched in SINEs, in particular Alu (Supplementary Figure 8) **d.** Relative distribution of STRs along AluJ (red), AluS (green) and AluY (blue). The stacked barplot shows the fraction of STRs corresponding to the linker (relative position *<* 0.6) or the polyA tail (relative position *>* 0.6) of AluJ (red), AluS (green) and AluY (blue). Fractions for linker: 33,653/69,122 (AluJ), 25,524/69,122 (AluS), 9,936/69,122 (AluY), ; fractions for polyA: 105,683/616,944 (AluJ), 405,683/616,944 (AluS) ; 102,926/616,944 (AluY). Fractions were compared with Fisher’s exact tests (pvalue *<* 2.2e-16 in all tests). Note that STRs located in Alu linker or polyA correspond exclusively to (*T*) *_n_*. The enrichment of STRs at AluJ linker has previously been noticed and may be explained by an accumulation of STR-interrupting mutations in polyA STRs 94. **e.** Enrichment of AluJ and AluS in rSTRs. The stacked barplots show the fraction of rSTRs or non-rSTRs located in AluJ, AluS, AluY and other RepeatMasker repeats, or located outside RepeatMasker repeats. rSTRs fractions: 796/8,401 (AluJ), 2,756/8,401 (AluS), 646/8,401 (AluY), 2,328/8,401 (other repeat), 1,875/8,401 (no repeat). non-rSTRs fractions: 1,052/13,967 (AluJ), 3,344/13,967 (AluS), 938/13,967 (AluY), 4,915/13,967 (other repeat), 3,718/13,967 (no repeat). Fractions were compared using Fisher’s exact tests ; pvalue = 4e-7 (AluJ), *<* 2.2e-16 (AluS), 6e-3 (AluY), *<* 2.2e-16 (other repeat), 5e-13 (no repeat).

### A complex transcriptional interplay between STR-initiating RNAs and Alu elements

We also investigated possible associations between rSTRs and Alu elements. The association between Alus and STRs has long been known ^61–63^ and Alu repeats have indeed been implicated in distal regulations ^64–66^. We confirmed that STRs are frequently found in short interspersed elements (SINEs) and Alu elements (Figure 6C and Supplementary Figure S8), where they correspond to Alu linker (A-rich sequence linking the two non-identical 7SL-derived monomers) and polyA tail (Figure 6D). This particular distribution is due to the fact that the HipSTR catalog is overrun by A/T-rich STRs ^17, 67^. We then studied the potential interplay between STR-initiating RNAs, rSTRs and Alu elements. Alu repeats are classified into three subfamilies with decreasing evolutionary ages: AluJ, AluS and AluY ^68^. We found that STRs involved in rSTRs are enriched in Alus, especially in ancient AluJ and AluS (Figure 6E), which are enriched in enhancer features ^66^. We confirmed that old Alus are enriched in enhancers using FANTOM5 enhancer catalog ^69^ (Supplementary Figure S9). The rSTRs were more enriched in Alu repeats than the eSTRs (4,276/8,458 vs. 6,094/22,037, odds ratio *→* 2.7, one-sided Fisher’s exact test p-value *<* 2.2e-16), suggesting a relationship between STR-initiating RNAs and Alu elements. We indeed noticed that the CAGE signal associated with Alu-antisense and STR-associated transcription was more elevated in AluJ, followed by AluS, and finally, AluY (Figure 7A). No difference was observed when looking at STR CAGE signal in the sense orientation of Alu elements (Figure 7A). Similar results were obtained using the classification made by Zhang *et al.* ^68^, which separates expressed Alu elements into five groups, B0–B4, depending on their presence in five primate genomes (human, chimpanzee, gorilla, orangutan, or macaque): the CAGE signal associated with STRs located in antisense orientation of Alu was lower in human-specific (i.e. B4) Alu repeats (Figure 7B), even when restricting the analyses to AluY (Supplementary Figure S10). This difference was not observed with STR-associated CAGE signal in Alu sense orientation (Figure 7B). These observations suggested that transcription of STRs located the antisense orientation of Alu elements is related to their evolution. Using TT-seq data ^70^, we showed that AluJ are more transcribed in the antisense orientation than AluS and AluY (Figure 7C). The AluY were previously shown to be expressed at lower rates than the older AluJ and AluS ^68, 71^. Together with our findings, this raised the possibility of an interplay between the transcription of Alu repeats and that of STRs located in Alu antisense orientation. The transcription of Alu elements was shown to be sensitive to *ω*-amanitine and it was hypothesized that Alu RNAs could be partly transcribed by Pol II ^71^. However, given the potential role of Pol II in regulating Pol III ^72^, we hypothesized that blocking Pol II transcription at STRs impairs Pol III-mediated Alu transcription. To test this hypothesis, we revisited the data generated by Baar *et al.* ^71^ and showed that the fractions of AluJ and AluS modulated by *ω*-amanitine treatment (i.e. adjusted p-value associated to fold change *<* 0.01) are greater than the fraction of modulated AluY (Figure 7D). Moreover, we used CFC-seq data ^50^ and showed that non-Alu Pol III genes do harbor TSSs in their 5’end (Figure 7E), suggesting that cap-trapping can capture Pol III RNAs and/or that some of these RNAs are indeed transcribed by Pol II. However, this CFC-seq pattern was not observed in the case of Alu repeats (Figure 7E and Supplementary Figure S11), suggesting that Alu transcripts are mostly transcribed by Pol III. We observed more CFC-seq TSSs at Alu 3’ends with more TSSs in the antisense orientation (Figure 7E and Supplementary Figure S11). Using a Poisson-binomial test, we found that these antisense TSSs have *→*5 more chance to be located within or close an STR (*<* 5bp) than expected by chance (see Methods section). The distribution of the CFC-seq TSSs along AluJ and AluS differs from that along AluY, with more TSSs located at the 3’ends of AluJ and AluS (Supplementary Figures S12 and S13). Conversely, we noticed a similar enrichment of Alu (Supplementary Figure S14) and (*A*)*_n_*/(*T*) *_n_* STRs at TSSs (Supplementary Figure S15). In agreement with the predictions of our MNNs, the ZNF384 motif was previously found enriched at Pol III genes ^73^ and expressed Alu elements ^68^. Leveraging Remap data, we found 5,181 ZNF384 peaks intersecting Alu genes (n = 49,308) but only 526 intersecting non-Alu Pol III genes (n = 12,008) indicating a clear enrichment of ZNF384 at Alu genes compared to other Pol III genes (Fisher’s exact test p-value *<* 2.2e-16, odds ratio = 2.6). We also observed that ZNF384 binds more frequently the core of AluJ than that of AluS and AluY (Supplementary Figure S16). We tried to leverage ZNF384 KD experiments to measure Alu modulations but, as previously reported ^74^, the RNA-seq signal at Alu elements was too low to find significant modulations. Overall, we concluded that Pol III transcription of Alu RNAs is linked to Pol II antisense transcription initiation at STRs.

**Figure 7.**
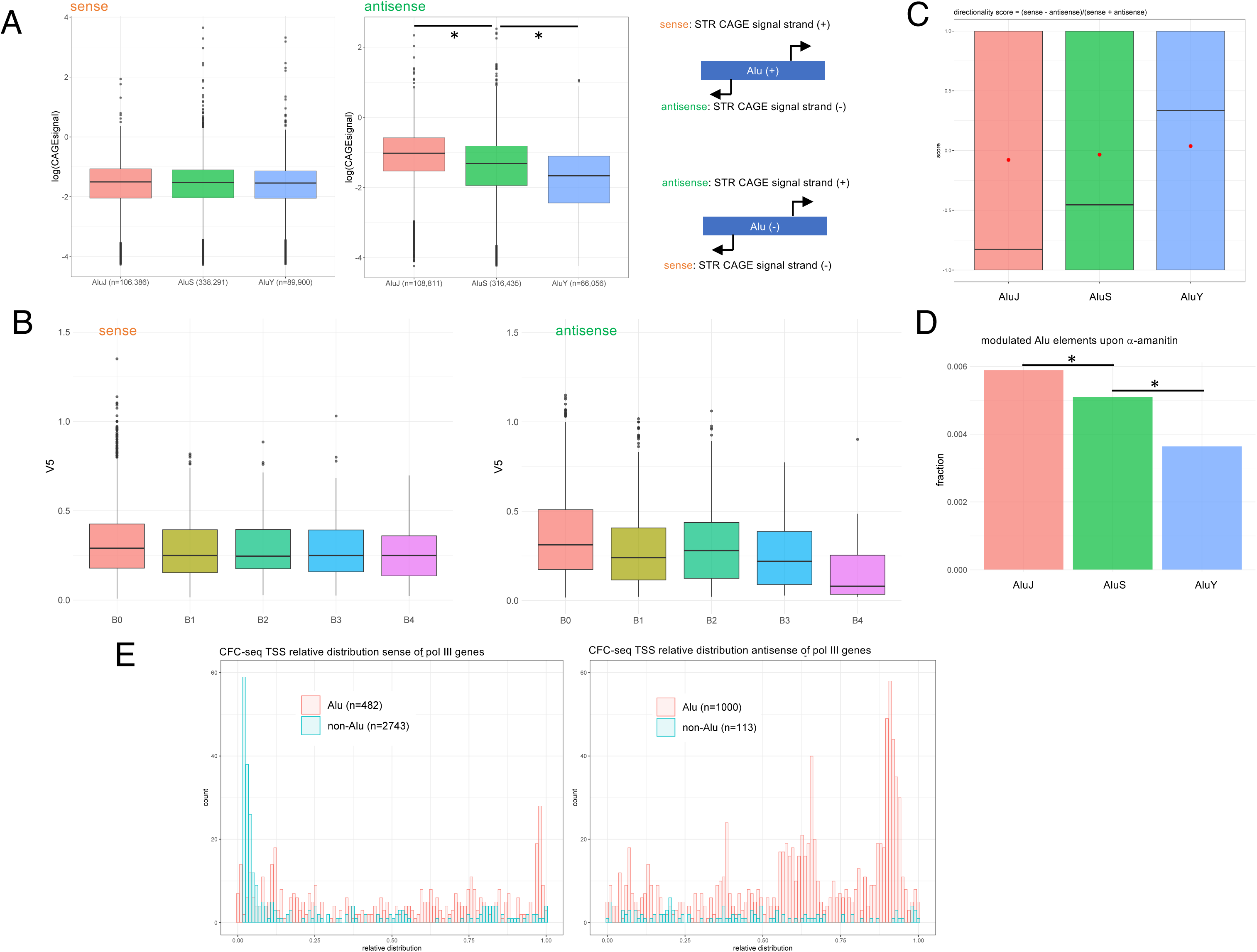
A complex transcriptional interplay between STRs and Alu elements. **a.** CAGE signal associated with STRs located in the sense (right) or antisense (left) orientation of AluJ (red), AluS (green) and AluY (blue). A diagram shows which CAGE signal is considered depending on the Alu strand (right). *, Wilcoxon test p-values *<* 2.2e-16. **b.** CAGE signal associated with STRs located in the sense (right) or antisense (left) orientation of expressed primate-specific Alu elements as defined in 68. The CAGE signal considered for each STR is similar to A. Here, expressed Alu elements were classified into five groups, B0–B4, depending on whether they were in four other primate genomes (chimpanzee, gorilla, orangutan, or macaque), with B4 defining the human-specific Alu elements (n = 103) ; B3 regrouping Alu present in human and chimpanzee, gorilla, orangutan (n = 126) ; B2 regrouping Alu present in human, chimpanzee and gorilla (n = 186) ; B1 regrouping Alu present in human, chimpanzee, gorilla, orangutan (n = 1,260) and B0 designating Alu present in human, chimpanzee, gorilla, orangutan and macaque (n = 12,286). Number of transcribed STRs located in the sense orientation = 89 (for B4), 76 (for B3), 128 (for B2), 826 (for B1) and 6,833 (for B0). Number of transcribed STRs located in the antisense orientation = 32 (for B4), 67 (for B3), 111 (for B2), 702 (for B1) and 6,478 (for B0). Wilcoxon test p-value = 0.56 for CAGE signals associated with STRs located in the sense orientation of Alu B4 and B3. Wilcoxon test p-value = 5e-3 for CAGE signals associated with STRs located in the antisense orientation of Alu B4 and B3. **c.** Directionality scores at AluJ (red, n=22,313), AluS (green, n=75,284) and AluY (blue, n=15,567) based on TT-seq data 70. Pairwise Wilcoxon tests were used to compare these scores in the three Alu classes (p-values *<* 2.2e-16 in all tests). Red dots represent the mean scores. **d.** Fraction of AluJ (red, 804/136,306), AluS (green, 1,673/327,453) and AluY (blue, 267/73,195) whose expression is significantly modulated (p-value *<* 0.01) by *ω*-amanitin treatment. Data are from 71 (GSE185485). Fractions were compared using Fisher’s exact tests (black lines). *, p-values *<* 1e-3. **e.** Relative distribution of CFC-seq TSSs in the sense (left) and antisense (right) orientation of Alu genes (red) and non-Alu pol-III genes (blue).

## Discussion

To study the synthesis of STR-initiating RNAs and assess their regulatory impact on gene expression, we developed fully-interpretable deep learning models, called MNNs, able to predict their expression level. Combining the output of independent deep learning models has been proposed in the past ^75^ and already used in genomics ^76, 77^. However, one key innovation of MNNs is that modules are not independent but learned one after another and additional modules are kept only if they increase the model accuracy. Besides, MNN hyper-parameters (learning rate, batch-size and filter size) are tuned for each block during training. There are several other differences with existing methods: (i) MNNs learn the filter size in each module while ExplaiNN uses a fixed filter size of 19bp ^76^. (ii) Similar to ExplaiNN, each module consists of one convolutional layer with one single filter. However, in ExplaiNN, the convolution is followed by exponential activation, two fully connected layers and dropout, not a ReLU and a single regression layer, which make it complicated to know where the filter is activated. Hence, while both ExplaiNN and MNN can identify which filter(s) is/are important, MNN can additionally identify subsequences wherein these filters are activated. (iii) the tiSFM method ^77^ uses the modular strategy and TF PWMs as filters in each convolution. However, filters can only be built from known motifs, which limits biological discovery and implications of other types of motifs ^9, 78^. (iii) Finally, MNNs contain much less parameters (*→* 1,000 parameters) compared to ExplaiNN and tiSFM, making them trainable with limited ressources. MNNs still have inherent limitations : (i) Because they use linear regression with the outputs of each module as variables, MNNs do not take into account motif interactions (e.g. synergism). This point could be explored adding new variables such as module output products for instance. (ii) MNNs use ReLU to highlight important regions of the input sequences. This might be improved using exponential activation, which has been shown to improve robustness and interpretability of learned representations in convolutional filters ^79^. (iii) At this stage, filter analyses are limited by HOMER identity score to JASPAR PWMs. Other resources could be used to take into account other motifs ^80^ and/or the diversity of TF binding mechanisms ^78^. For example, the Codebook consortium ^81^ has recently published new PWMs for several TFs, including one for ZNF362 that is similar to that of ZNF384 ^49^, making it another candidate regulator of STR transcription.

The predictions of our MNN models were used to compute rSTRs that take into account the impact of SNPs located around STRs, not STR length variations. The CAGE signal detected at STRs did not appear directly linked to their length (Supplementary Figure S1) and, leveraging CFC-seq data, we could not find difference in terms of both CAGE signal and STR length between rSTRs and non-rSTRs (Figure 5C and D). Still we cannot exclude that STR length variation influences the impact of SNVs, as previously reported ^25^, and thereby STR-initiating transcription. Ideally, one would need a measure of transcription initiation that is less sensitive to mapping issues in repeated sequences. In addition, our models were trained on the reference genome, and their predictions could probably be improved by training on the genome of cells where expression has been measured ^82, 83^. Hence, combining cap-trapping and long read sequencing ^17, 50, 84, 85^ with whole genome sequencing, at a large scale, could generate appropriate data and allow more thorough computations of combined e/rSTRs that would take into account both STR length and surrounding SNPs.

Note that SNPs mutate at a rate on the order of 2.0-2.5 *↑*10*^→^*8 mutations per nucleotide position per generation, while a mutation rate of *→*1.5*↑*10*^→^*3 per STR per generation is observed for dinucleotide repeat polymorphisms ^23^. This implies that the variance of STR length, previously used to compute eSTRs ^34^, is greater than that of the predictions made by MNNs, which are based on the presence of SNPs, and which has been used to compute our rSTR catalog. This point might explain why we detected less associations (n=14,340) than Fotsing *et al.* (n=28,375), who used STR length variations ^34^.

The rSTRs computed here revealed a complex interplay between the transcription of Alu elements and that of STRs in their antisense orientation, opening up potential new avenues of research. The observation that antisense Pol II transcription was lower in recently inserted Alu, i.e. active AluY and human-specific Alu, suggested that this transcription appeared after retrotransposition, presumably through *de novo* STR mutations ^19, 62, 86^, which created transcription initiation sites and concomitantly abolished the ability to retrotranspose. Alu-antisense transcription at STRs could be implicated in the control of distal regulations orchestrated by old and conserved Alu elements, for instance by controlling the synthesis of regulatory RNAs and/or by acting as RNA sponge through double-stranded RNA formation to prevent target recognition ^64^. Finally, given the potential role of ZNF384 in transcription at STRs, it would also be interesting to study Alu expression where the activity of this TF is altered, for example in acute lymphoblastic leukemia ^87^ and mixed phenotype acute leukaemia ^88^.

## Methods

### Modular Neural Network

The MNN models were implemented in PyTorch. They use as input 101bp long sequences centered around STR 3’ end (as defined by the strand of the transcription). The CAGE signal correspond to raw tag count along the genome with a 1bp binning and Q3 quality mapping filter. At each position of the genome, the mean tag count across 988 human librairies was computed. The values obtained at each position of a window encompassing the STR*±* 5bp were then summed and normalized (i.e. divided by the STR length + 10 bp) to limit the impact of CAGE noise signal observed along STRs as in ^17^. The predicted variable correspond to the rank of the CAGE signal. Model performances were evaluated computing Spearman correlation between predicted and observed ranks.

Each MNN module hyperparameters are determined using an Asynchronous Successive Halving Algorithm (ASHA) ^89^ implemented in RayTune (https://docs.ray.io/en/latest/tune/api docs/schedulers.html). Briefly, ASHA considers each hyperparameters combination in the search space and aggressively terminates unsuccessful hyper-parameter combinations to allocate more budget towards better performing combinations. This is particularly effective when considering large training times and small search spaces ^89^, which is the case for MNNs as each module search space is composed of a handful of parameters (convolution filter size, momentum and learning rate). Each hyperparameter combination is then associated with a MNN and trained using the ADAM optimizer ^90^ with performance computed on an held-out validation set. After the best configuration has been found, the module is frozen and another module is added to the MNN which hyperparameters are yet to be determined by ASHA.

Finally, the validation performance on held-out data is kept in memory after each time a new module is added. Then, if performance does not improve according to a pre-defined threshold (here 0.005) after adding a new module, the model is given 5 tries to improve. If the model does not improve, training is terminated and the final version of the model is returned.

The source code of the model, alongside scripts and Jupyter notebooks are available at https://gite.lirmm.fr/ml4reggen/rstr.

### GTEx long read RNA sequencing analyses

GTEx long reads data were downloaded from GTEx portal (see ‘Data Availability’). To find long reads initiating at STRs, we looked for GENCODE V26 transcript start sites in a window of 5bp around STRs using the following bedtools window command line:

windowBed -w 5 -a gencodev26.transcript.start.sites.bed -b hg38.hipstr\

_reference.cage.bed

This list of 2,970 (STR, long read) pairs was further used to compute the correlation between STR length and expression of associated long reads for each of the GTEx samples. The distribution of the correlations are shown Supplementary Figure S1.

To assess the tissue-specific expression of GTEx long reads, we used Gini coefficient. Briefly, the Lorenz curve is a classical graphical representation of the income distribution between individuals in economics. It is obtained by ordering the individuals by their income, and by calculating the cumulative part of income in function of the cumulative part of individuals. In the case of an equal distribution between all individuals, the curve follows the line y = x. Otherwise, it is found under this line. The surface A is the area between the Lorenz curve and the line of perfect equality of distribution. The surface B is the area between the Lorenz curve and the perfect inequality curve (all the income belongs to a single individual). The Gini coefficient is defined by

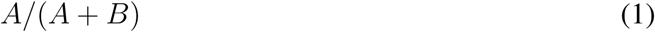

It is equal to 1 if all the incomes belong to a single individual and equal to 0 if the incomes are equally distributed. For long read expression, we computed the Gini coefficient of each transcript by ordering the samples by the transcript expression and by calculating the cumulative part of expression in function of the cumulative part of samples.

### Filter analyses and motif discovery

We downloaded the ‘latest’ versions of the motifs from the ‘CORE’ collection of the ‘Vertebrates’ group in Jaspar 2022 and the ‘latest’ versions of the motifs from the ‘POLII’ collection in Jaspar 2020 using the Jaspar REST API. These motifs were then merged in a single file in MEME format. We converted this file into Homer motif format using our in-house tool (pwm2homer.py). We used this file to search for motifs similar to *de novo* motifs when running Homer. For each class of STR model and each module, we created three fasta files: (i) The ‘positiveMnnHits.fasta’ file, which contains all sequences of the size of the CNN filter for the given module, starting at positions where the CNN returns a positive score (ReLU *>*0). (ii) The ‘negativeMnnHits.fasta’ file, which contains all sequences of the size of the CNN filter for the given module, starting at positions where the CNN returns a null or negative score and entirely included in the 101 nucleotide sequence. (iii) The ‘otherPositiveMnneHits.fasta’ file, which contains the longest sequences from the positive hits of all other modules of the given model. For each module, we ran Homer with the ‘positiveMnnHits.fasta’ file as the foreground and either the ‘negativeMnnHits.fasta’ (negHitBg) or ‘otherPositiveMnneHits.fasta’ (otherHitBg) file as the background. The length of *de novo* motifs (using the --len option) varies from 5 to the filter size when the filter size is greater than 5; otherwise, it is set to the filter size. We chose a size of 5 because it corresponds to the size of the smallest motif in our Jaspar motifs. The results from Homer were parsed and compiled into a csv table. The source code, alongside scripts, results and Jupyter notebooks are available at https://gite.lirmm.fr/ml4reggen/rstr.

### ZNF384 KD ENCODE experiments

The bam files for the ENCSR103URL (CRISPRi targeting ZNF384) and ENCSR016WFQ (control) experiments were downloaded from the ENCODE portal (see ‘Data Availability’). Reads were counted using featureCounts (subread v2.0.1) using the FANTOM gtf file combining ENCODE, FANTOM and CFC-seq long reads transcripts (table5pENSTpCAT.gtf) ^50^. Genes were filtered using the edgeR v4.0.16 package to retain only genes with sufficiently high counts for statistical analysis. Differential expression analysis was performed using DESeq2 v1.42.1. Multiple hypothesis adjusted p-values were calculated using the Benjamini-Hochberg procedure to control for false discovery rate (FDR). The logFC of ZN384 (ENSG00000126746.18) was −0.69 with an adjusted p-value of 7.5e-4 (Supplementary Table S2). Genes with at least one transcript 5’end located in a window of 5bp of an STR were considered as genes with transcripts initiating at STRs (n = 41,766 out of 53,566, Supplementary Table S2), which represent the majority of the genes (78%).

### rSTR computation

We associated STRs and genes using the bedtools window ^91^ and the following command line:

windowBed -w 100000 -a gencodeV19.gene_coord.bed -b hg19.hipstr_reference.cage.bed

We then intersected the STR 3’end +/− 50 bp coordinates with the VCF file from GTEx to retrieve the individual SNPs within said window, introduced the SNPs in the STR reference sequences and predicted their MNN score using a custom-made python script. Finally, using MANTA, we computed the associations between each loci MNN scores and their associated gene. The source code of the rSTR computation procedure is available at https://gite.lirmm. fr/ml4reggen/rstr. GTEx data were obtained through dbGAP under phs000424.v7.p2. The rSTR computation procedure is available as a Nextflow workflow within the rstr repository and MANTA is available as a R package at https://cran.r-project.org/web/ packages/manta/index.html. As in ^34^, we restricted our computations to 17 tissues with at least 100 samples.

To compare newly computed rSTRs with published eQTLs ^33^, the bedtools intersect ^91^ was used to intersect STRs (50bp around 3’end according to strand of transcription) with SNP coordinates. Similar analyses were performed with the credible sets of causal variants listed in CAUSALdb ^55^. To compare rSTRs to previously published eSTRs ^34^, we consider the STR coordinates as defined in hg19 HipSTR catalog (see ‘Data Availability’). The link to download eQTLs, sQTLs and eSTRs are provided in the ‘Data Availability’ section.

### Evaluation of rSTR significance

To assess the presence of rSTRs in Pol II-mediated DNA-DNA interactions, we used ENCODE Pol II ChIA-PET data from 5 different cell lines: K562 (ENCFF511QFN), HepG2 (ENCFF360QPK), HCT116 (ENCFF322FOT), GM12878 (ENCFF040KUS), A549 (ENCFF946FGU). We considered rSTRs and rGenes implicated in significant associations and counted the total number of (STR, gene) pairs found in ChIA-PET reads. Specifically, we intersected each ChIA-PET fragment coordinates with that of rSTRs and rGenes and created a set of possible ChIA-PET-detected pairs. We then computed the fraction of these pairs corresponding to significant rSTR associations, excluding associations implicating rSTRs and rGenes located within the same ChIA-PET fragment. The same analysis was repeated with all (tested STR, tested gene) pairs. These two fractions were compared using Fisher’s exact test. The odds ratio are shown Figure 4A.

To evaluate the presence of rSTRs in RNA-DNA interaction, we used RADICL-seq data from 19 different cell types (Lambolez*et al.*, in preparation). We first intersected the coordinates of RADICL-seq read RNA portions with coordinates of transcripts (ENCODE or FANTOM) and/or RNAs detected by cap-trapped long read ONT RNA sequencing (CFC-seq) ^50^ initiating at rSTRs (i.e. whose 5’ end is located in a window of 5bp around rSTRs). We intersected the coordinates of RADICL-seq read DNA portions with coordinates of rGenes. As for ChIAPET, we counted the number of (rSTR, rGene) pairs detected by RADICL-seq. For comparison, we defined a set of non-rSTRs as among the tested pairs those implicating STRs never found associated with the expression of any gene and genes whose expression was never found associated with STR transcription (n = 36,199). The fraction of rSTRs detected in RADICL-seq and that of non-rSTRs were compared using Fisher’s exact test. Note that comparing fraction of rSTRs detected in RADICL-seq to that obtained with all tested rSTRs yielded non significant results, likely due to the presence of false negative rSTRs ^54^.

To compute enrichment of CAUSALdb variants in rSTRs, we intersected the 101bp-long regions centered around the 3’end of STRs engaged in rSTRs (n=8,458) and all tested STRs (n=22,491) with coordinates indicated in CAUSALdb ^55^. The distribution of CAUSALdb variants around STRs was computed by intersecting variant coordinates with regions encompassing 100bp located upstream STR 5’ends or 100bp downstream STR 3’ends (variants located within STRs were not considered in this analysis). The number of variants at each position of these regions was computed and shown Figure 5A.

The CAUSALdb variants located 100bp up/downstream STRs were annotated with RADAR using radar.py ^56^. RNAs initiating at STRs engaged or not in rSTRs were identified by intersecting the coordinates of CFC-seq long read 5’ends ^50^ with STR coordinates. The coordinates of these RNAs were then intersected with those of CLIP peaks listed in POSTAR3 CLIPdb ^57^ using bedtools intersect ^91^ option -s. The relative position of the CLIP peaks with transcripts were computed as, if transcript strand=”+”

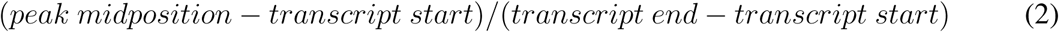

or otherwise

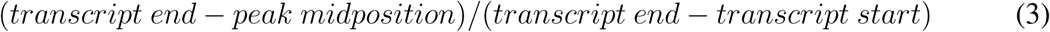

The gene coordinates were retrieved using ENSEMBL Biomart. The distances between STR and gene in rSTRs were computed no matter the strand of the STR or the gene and set to 0 if the STR was located within the rGene (STR start *>* rGene start and STR end *<* rGene end), to the difference between the rGene start and the STR end if the STR was located upstream the rGene (STR end *<* rGene start), to the difference between the STR start and the rGene end if the STR was located downstream the rGene (STR start *>* rGene end). The distance between STRs and their closest genes was computed using bedtools closest ^91^ using the option -D:

closestBed -a str_coordinates -b gene_coord.bed -D a

### MNN score and gene expression correlations

MANTA does not provide correlation coefficients between MNN score and gene expression, only p-values. To generate Table S7, we collected, for each rSTR and for each individual, the MNN score on both STR alleles and computed the average between those two scores. We then computed, for each rSTR, the Spearman correlation between vector of MNN scores computed in all GTEx individuals and that of gene TPMs in the same individuals using the Python *scipy.stats.spearmanr*. For each rSTR, all tissues have been considered (i.e. the length of the vectors used to compute correlation can differ from one association to another, in contrast to rSTR computations).

### RepeatMasker and Alu analyses

The coordinates of all human repeated sequences were retrieved from UCSC RepeatMasker track and the coordinates of Alu elements were intersected with those of STRs engaged in rSTRs or eSTRs using the bedtools intersect ^91^ with options -f 1 (to make sure 100% of the STR is located within Alu) and, when required, -s and -S. Similarly, the Supplementary Table S4 of ^68^ was converted into bed file using awk command line and intersected with the coordinates of STRs. TT-seq data are from ^70^. The directionality score was computed intersecting the binned signal (GSE75792) with Alu coordinates and following the formula for each Alu element:

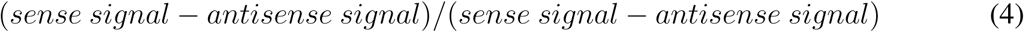

To generate Supplementary Figure S9, the coordinates of FANTOM5 enhancers were intersected with that of Alu repeats in RepeatMasker. We then counted the number of AluJ, AluS and AluY using grep option -c. For comparison (see legend of Supplementary Figure S9), we generated 63,285 random sequences using the following command line:

bedtools shuffle -i F5.hg38.enhancers.bed -g hg38.chrom.sizes -chrom -excl F5. hg38.enhancers.bed

The *ω*-amanitin data are from Baar *et al.* ^71^. Alu and non-Alu Pol III genes coordinates were retrieved from Pol3Base ^92^. The relative distribution of CFC-seq TSSs (5’end position) along Alu and other Pol III genes was computed using the bedtools intersect ^91^ with -s (for sense, Figure 7 left panel) and -S (for antisense, Figure 7 right panel) options as, if gene strand=”+”

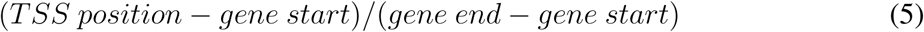

or otherwise

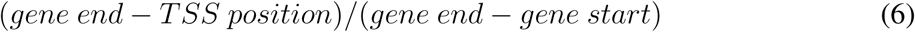

The enrichment of CFC-seq TSSs (i.e. long read 5’end) in Alu STRs was computed restricting the analyses to Alu containing at least an STR and at least one TSS (number of trials) and using a Poisson binomial test: for each Alu element, the hypothesized probability of success was computed as the ratio between the length of the STR and the length of the Alu element. Number of success corresponded to the number of Alu elements where a TSS is located within or close an STR (using bedtools window -w 5 ^91^). A p-value was computed using the Chernoff bound https://en.wikipedia.org/wiki/Poisson_binomial_distribution. The ratio between the total number of success and the sum of prior probabilities (i.e. expectation) provided an odds ratio (*→* 5 in that case). The relative distribution of Alu (Supplementary Figure S14) and (*A*)*_n_*/(*T*) *_n_* STRs (Supplementary Figure S15) around TSSs was computed using bedtools window -w 2000 ^91^).

### Other bioinformatic analyses

When required, hg19 coordinates were liftovered to hg38 using the UCSC liftover command line tool ^93^ and the hg19ToHg38.over.chain file available at this url https://hgdownload.soe.ucsc.edu/gbdb/hg19/liftOver/hg19ToHg38.over. chain.gz.

All statistical tests were performed with R (*wilcoxon.test*, *fisher.test*) or Python (*scipy.stats.mannwhitneyu*) as indicated. When indicated, p-values were corrected for multiple testing using R *p.adjust* (method=”BH”).

### Data Availability

- HipSTR: https://github.com/HipSTR-Tool/HipSTR-references/raw/master/ human/hg19.hipstr_reference.bed.gz
- GENCODE V19: https://ftp.ebi.ac.uk/pub/databases/gencode/Gencode_human/release_19/gencode.v19.annotation.gtf.gz
- GENCODE V26: https://ftp.ebi.ac.uk/pub/databases/gencode/Gencode_human/release_26/gencode.v26.annotation.gtf.gz
- GTEx: long reads https://gtexportal.org/home/downloads/adult\protect\discretionary{\char\hyphenchar\font}{}{}gtex/long_read_data/quantificatgencode.tpm.txt.gz
- CAGE signal at STRs: https://gite.lirmm.fr/ibc/deepSTR/-/blob/master/Data/hg19.hipstr_reference.cage.bed?ref_type=heads
- GTEx V7 data are available with controlled access through dbGaP (phs000424.v7.p2).
- eQTLs and sQTLs: https://gtexportal.org/home/downloads/adult-gtex/qtl
- eSTR coordinates were retrieved from Supplementary Table 1 of ^34^ https://static-content.springer.com/esm/art%3A10.1038%2Fs41588-019-0521-9/MediaObjects/41588_2019_521_MOESM3_ESM.xlsx
- HOMER motif discovery tool: http://homer.ucsd.edu/homer/
- JASPAR: The ‘latest’ versions of the motifs from the ‘CORE’ collection of the ‘Vertebrates’ group in Jaspar 2022 and the ‘latest’ versions of the motifs from the ‘POLII’ collection in Jaspar 2020 were downloaded using the Jaspar REST API.
- Remap2022 and ZNF384 ChIP-seq: https://remap2022.univ-amu.fr/
- ZNF384 KD: ENCSR103URL (CRISPRi targeting ZNF384) and ENCSR016WFQ (control) were downloaded from ENCODE portal https://www.encodeproject.org/.
- ENCODE Pol II ChIA-PET data from 5 different cell lines, namely K562 (ENCFF511QFN), HepG2 (ENCFF360QPK), HCT116 (ENCFF322FOT), GM12878 (ENCFF040KUS) and A549 (ENCFF946FGU), were downloaded from ENCODE portal https://www.encodeproject.org/.
- CFC-seq XXX
- RADICL-seq XXX
- The coordinates of ENCODE Candidate Cis-Regulatory Elements (cCREs) combined from all cell types were retrieved from UCSC Table browser (https://genome.ucsc.edu/cgi-bin/hgTables, group: ‘Regulation’, track: ‘ENCODE cCREs’).
- CAUSALdb: https://onedrive.live.com/?authkey=%21AAkjBQGnsKq5BFU&id=F7BB08E3479FE7EA%21331&cid=F7BB08E3479FE7EA
- RADAR: https://github.com/gersteinlab/RADAR/blob/master/
- The coordinates of RepeatMasker sequences, including Alu elements, were retrieved from UCSC Table browser (https://genome.ucsc.edu/cgi-bin/hgTables, group: ‘Repeats’, track: ‘RepeatMasker’).
- Pol3Base: https://rna.sysu.edu.cn/pol3base/download.php
- POSTAR3 CLIPdb: http://111.198.139.65/RBP.html
- FANTOM5 enhancers: https://fantom.gsc.riken.jp/5/datafiles/reprocessed/hg38_latest/extra/enhancer/F5.hg38.enhancers.bed.gz
- TT-seq: https://www.ncbi.nlm.nih.gov/geo/download/?acc=GSE75792&format=file&file=GSE75792%5Fbinning%5Fanno%2Etxt%2Egz
- *ω*-amanitin treatment: https://www.ncbi.nlm.nih.gov/geo/download/?acc=GSE185485&format=file&file=GSE185485%5Fdifexp%2Egtf%2Egz
- Alu classification from Supplementary Table S4 of ^68^

### Code Availability

Source code of MNN models, detailed results of filter analyses, a readme.txt file and other instructions for installing and running rSTR computations are available on our git repository https://gite.lirmm.fr/ml4reggen/rstr.

## Acknowledgements

We thank all members of the IGMM/LIRMM/IMAG Computational Regulatory Genomics team and of the FANTOM consortium for insightful discussions and/or critical readings of the manuscript. We are indebted to the researchers around the globe who generated experimental data used in this work and made them freely available. This project has received financial support from the CNRS through the MITI interdisciplinary and the International Associated Laboratory programs, as well as from the French National Research Agency (ANR*↓*22*↓*CE45*↓*0031*↓*01 and ANR*↓*22*↓*PESN*↓*0013). M.G. was supported by a *Conventions Industrielles de Formation par la Recherche* (CIFRE) PhD fellowship from SANOFI R&D.

## Author contributions

M.G., C.V., D. G-M., L.B., C.C., R.G. and C-H.L. conceived the scientific strategy, analyzed and interpreted data. M.G. developed MNN models. M.G. and A.V. evaluated MNN predictions. M.G. and D. G-M. computed rSTRs. M.G. and C-H.L. analyzed rSTR results. C.V., L.C., Q.B. and C-H.L. analyzed MNN filters. M.R. re-analyzed ZNF384 KD RNA-seq data. P.C., C.C., L.B., R. G. and C-H.L acquired fundings. M.G., C.V. and C-H.L. wrote the manuscript. All authors have read and approved the manuscript.

## Supplementary Figures

**Supplementary Figure S1.**
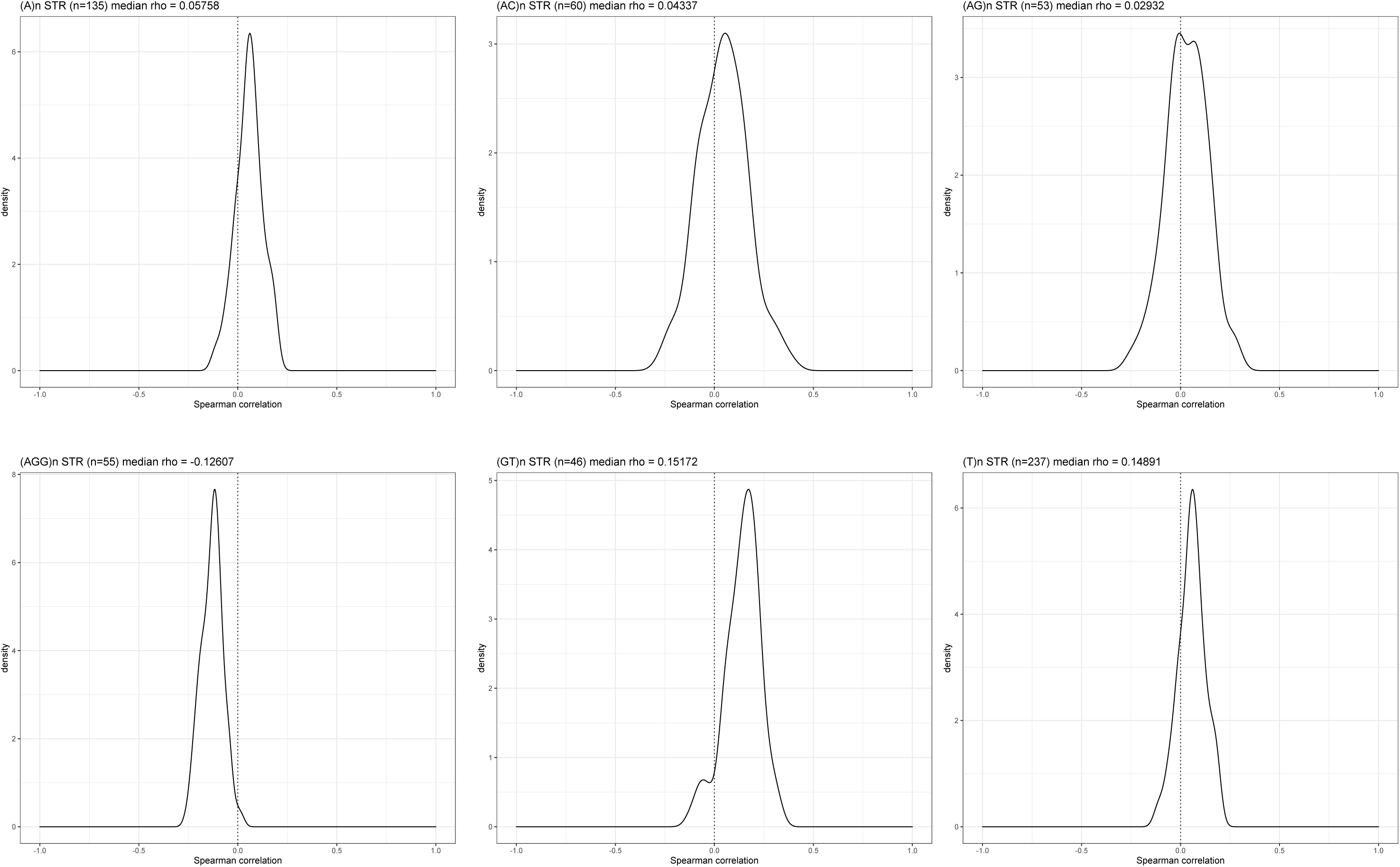
Spearman correlation between STR-initiating transcripts and STR length. Spearman correlation between the length of STRs and GTEx long reads TPM. The TPM of GENCODE V26 transcript whose start sites are located in a window of 5bp around STRs are considered. STRs are distinguished by the nature of the repeated k-mers (STR class). The distribution of the Spearman correlation coefficient rho between STR-initiating transcript expressions in each GTEx sample and STR lengths in the reference genome is shown (number of rho in each distribution = 92). The number of STRs considered and the median rho are indicated.

**Supplementary Figure S2.**
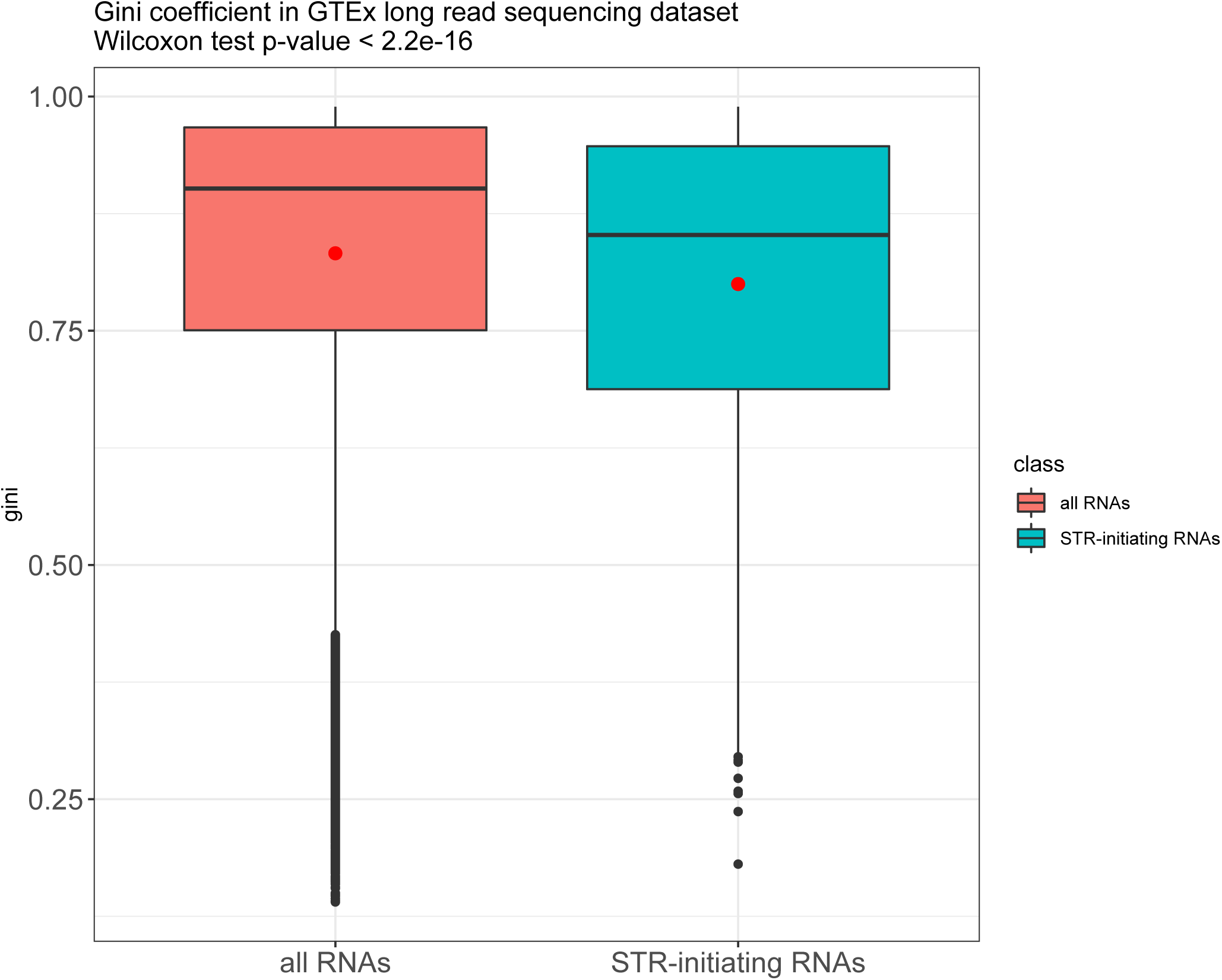
Gini coefficient of STR-initiating transcripts. The Gini coefficient (see Methods) of each GTEx transcript was computed to assess expression ubiquity in the 92 GTEx samples. The distribution of all coefficients is shown as a boxplot (red). The lower and upper hinges correspond to the first and third quartiles (the 25th and 75th percentiles). The upper whisker extends from the hinge to the largest value no further than 1.5×IQR from the hinge (where IQR is the interquartile range or distance between the first and third quartiles). The lower whisker extends from the hinge to the smallest value at most 1.5×IQR of the hinge. Data beyond the end of the whiskers are plotted individually. Red dots correspond to mean values. The red dot indicates mean coefficient. The blue boxplot shows the distribution of the Gini coefficient for 2,970 STR-initiating transcripts.

**Supplementary Figure S3.**
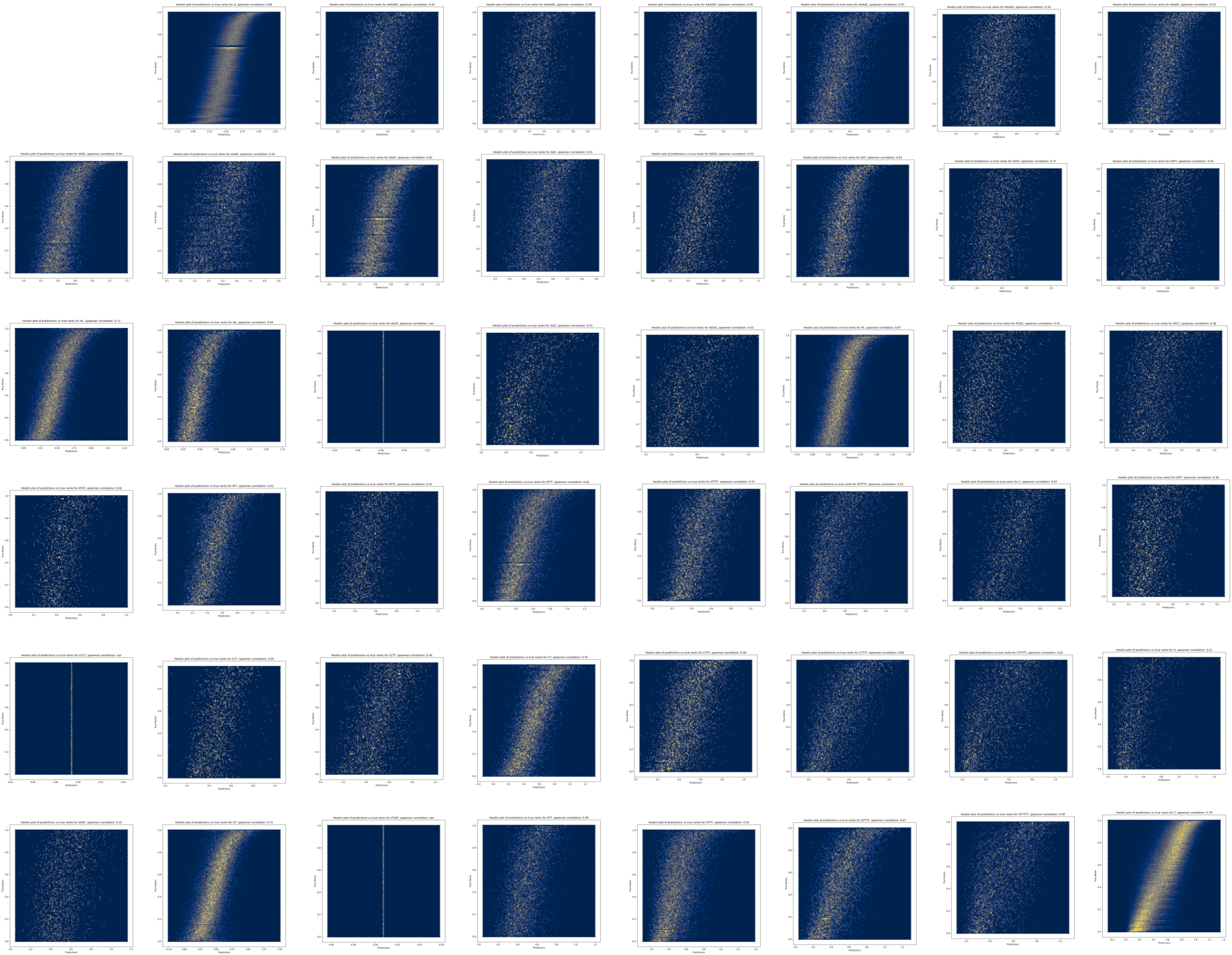
MNN performances. Hexbin plots showing predicted (y-axis) vs. observed ranks (x-axis) for different STR classes. The Spearman correlation coefficient is indicated on top of each panel.

**Supplementary Figure S4.**
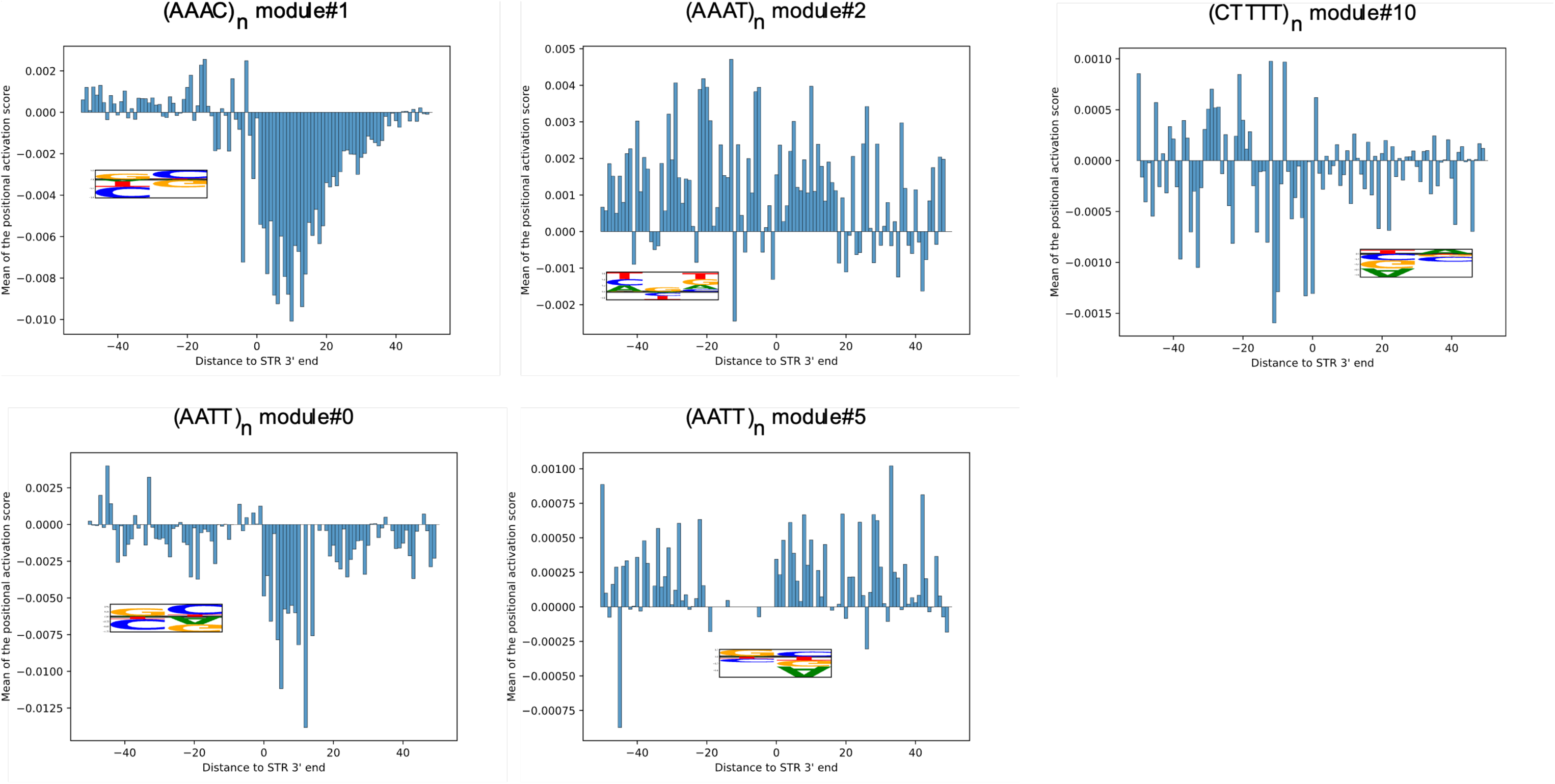
Examples of filters corresponding to short MNN filters that did not match a PWM. The distribution of filter activations around STR 3’end is shown. Activation corresponds to the multiplication of the mean of the activation score after the ReLU layer for all sequences of the model by the weights of the first regression layer and by the weight of the module in the second regression layer. The STR class and the number of the module are indicated on the top of the panel. Logo representations of the filters are shown.

**Supplementary Figure S5.**
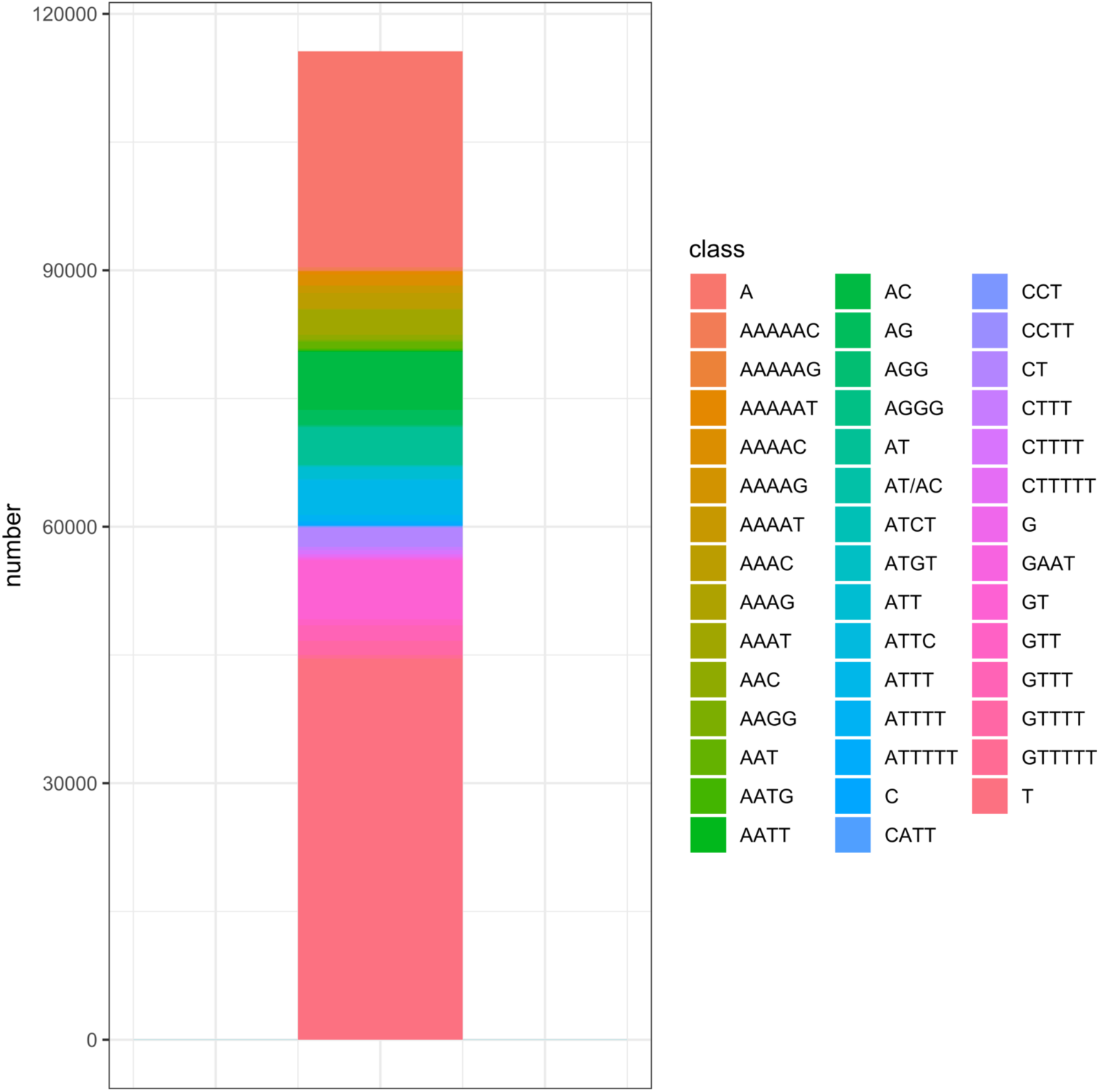
Distribution of STR classes considered for rSTR computations. y-axis, number of STRs in each STR class.

**Supplementary Figure S6.**
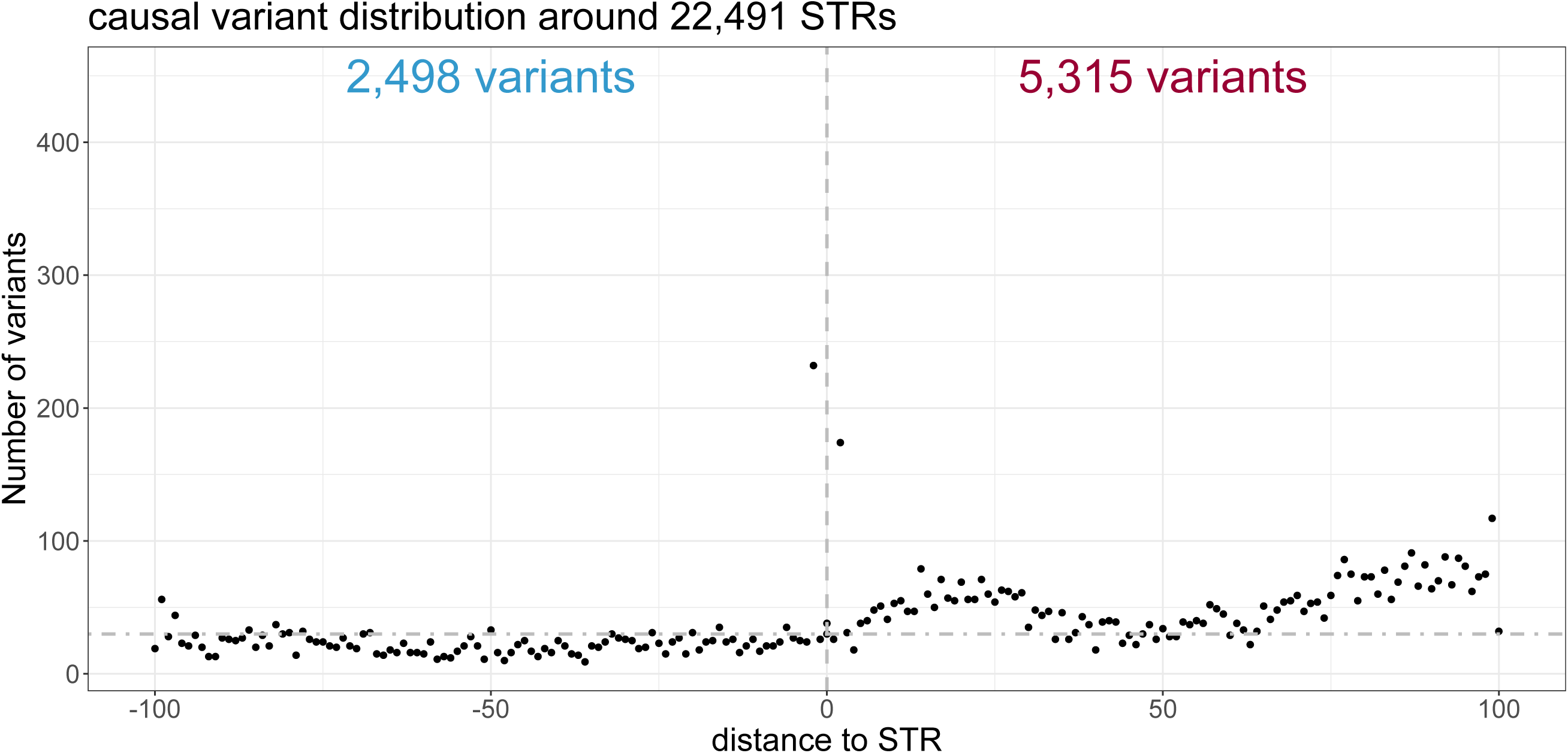
Distribution of causal variants around all tested STRs. STR sense is defined by the strand of the CAGE signal (right). Hence, one single STR associated with CAGE signal on both strand will have 2 downstream and 2 upstream sequences. x-axis: distance to STR 5’ or 3’ ends ; y-axis: number of variants (each dot is a variant). Vertical dashed grey line indicates the position of the STR. Horizontal dashed grey line is arbitrary drawn to highlight the difference between the number of variants located upstream and downstream STRs. number of variants located downstream = 5,315 ; number of variants located upstream = 2,498 ; Fisher’s exact test p-value < 2.2e-16, odds ratio ∼ 2.47

**Supplementary Figure S7.**
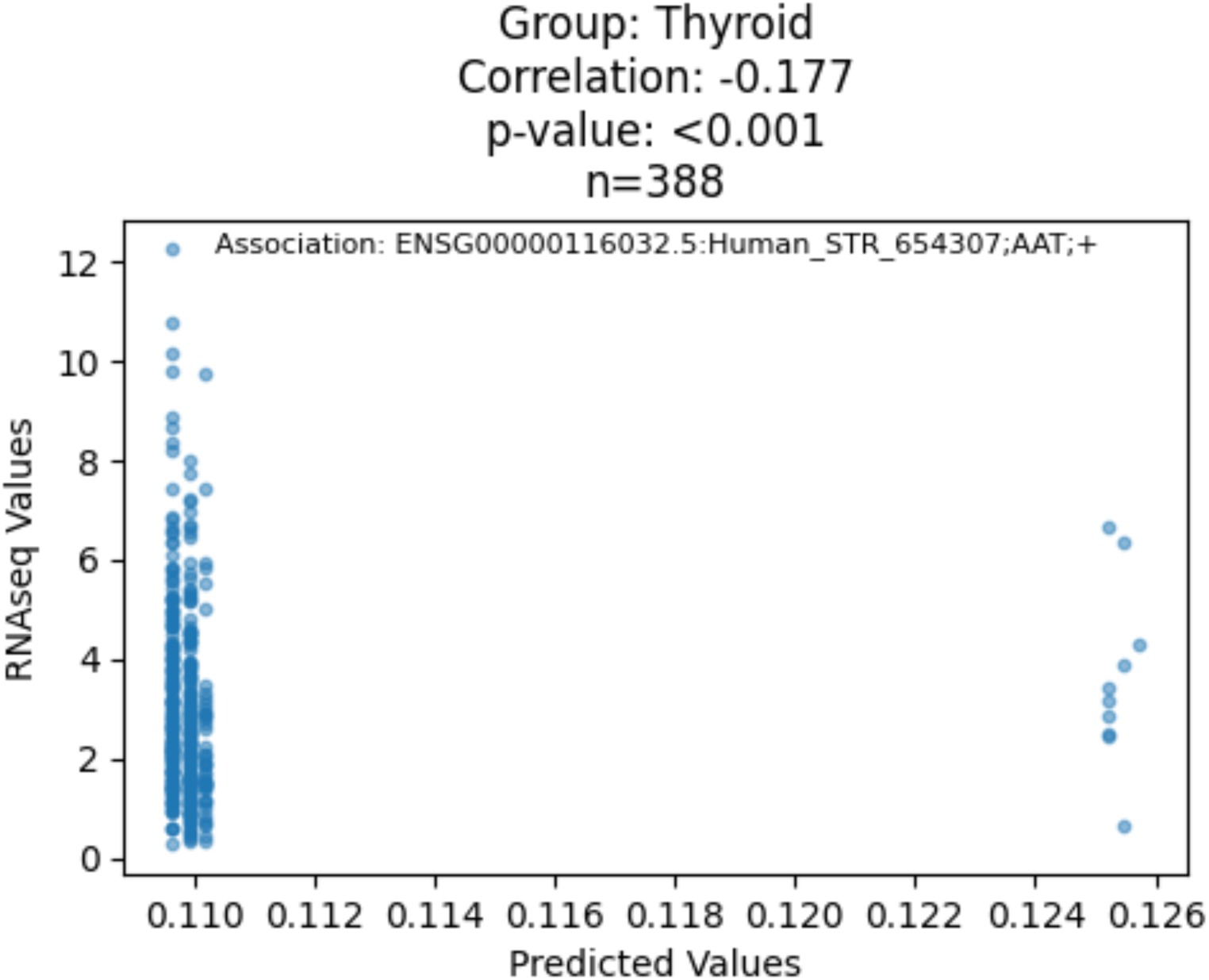
MNN predictions at Human_STR_654307;AAT;+ (x-axis) vs GRIN3B expression values (TPM, y-axis) in Thyroid tissue (see Supplementary Table S4A). rs12151021 is one of the variants considered for the prediction. The range of x values is small indicating that this variant has a limited - but significant – impact on GRIN3B expression. Covariates are not included in the correlation computation. All tissues available in GTEx V7 have been considered (i.e. no restriction to the 17 tissues used for rSTR computation). The Spearman correlation between MNN score and rGene expression is shown.

**Supplementary Figure S8.**
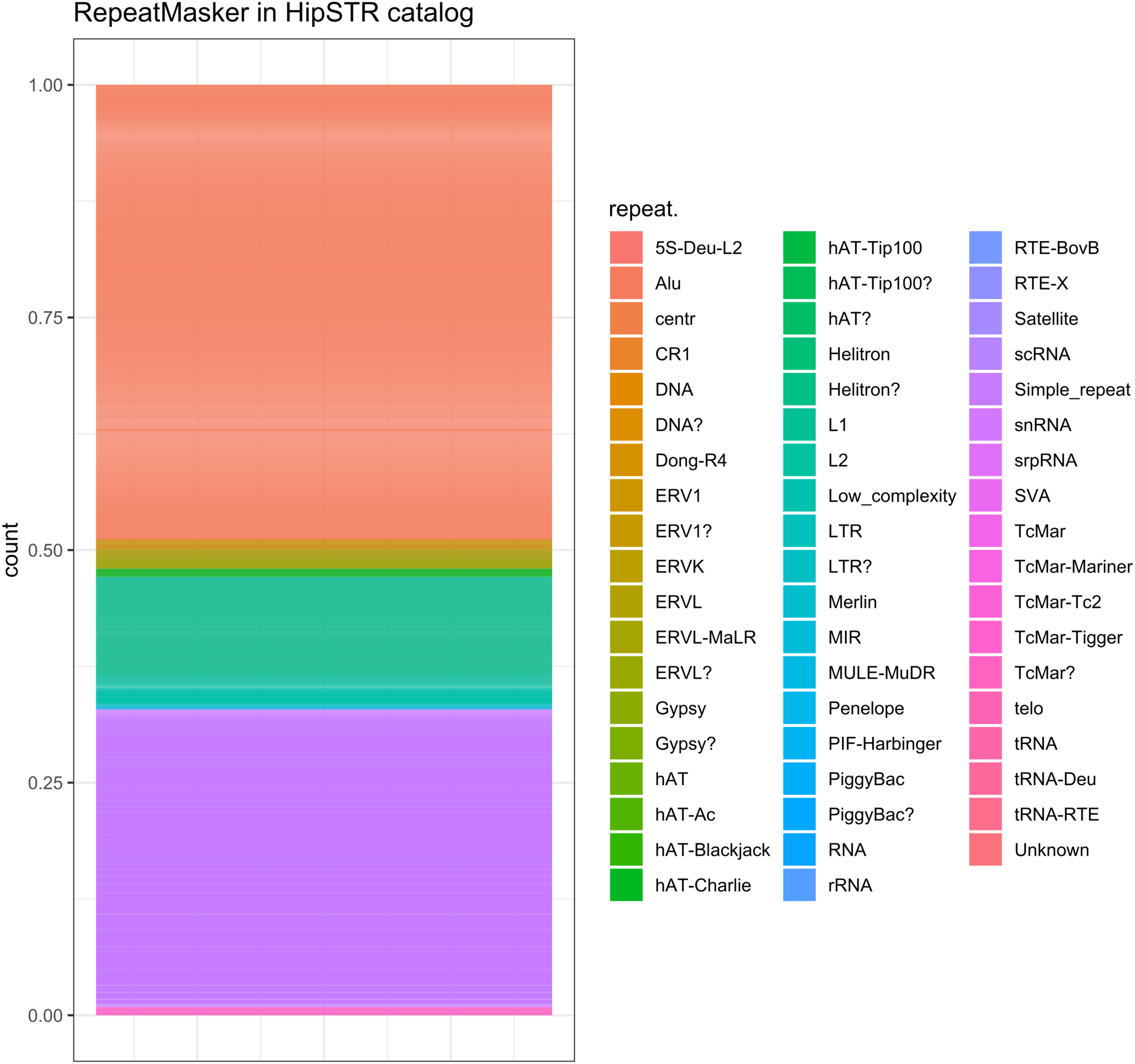
Distribution of HipSTR STRs in RepeatMasker families. y-axis, fraction of STRs in each RepeatMasker family.

**Supplementary Figure S9.**
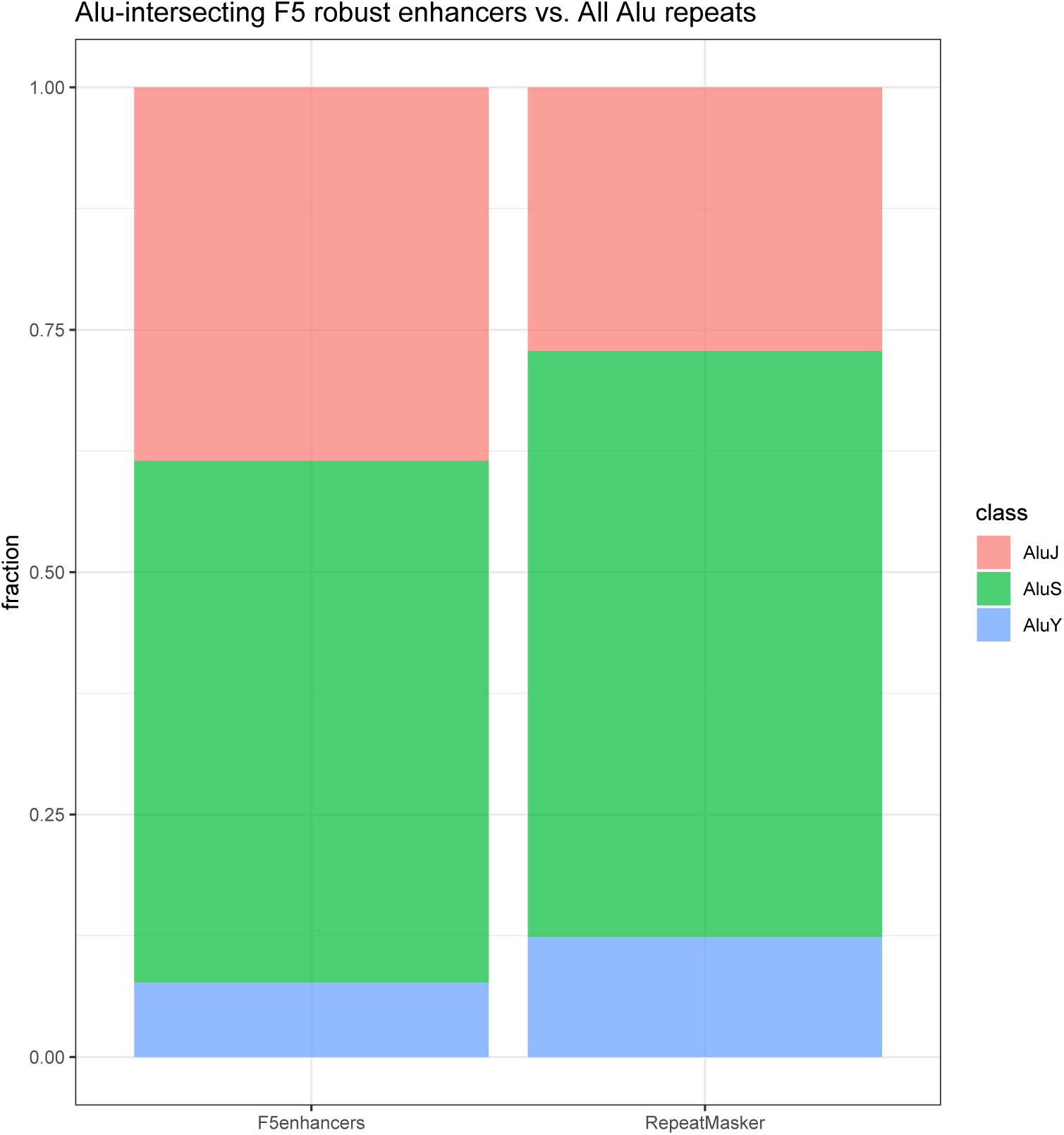
Fractions of Alu repeats in FANTOM robust enhancers. The coordinates of FANTOM5 enhancers were intersected with that of Alu repeats in RepeatMasker (n = 5,514). We then counted the number of AluJ (n = 2,115), AluS (n = 2,951) and AluY (n = 422). The fractions of AluJ (332,819/1,228,449), AluS (739,060/1,228,449) and AluY (151,627/1,228,449) present in RepeatMasker are shown for comparison. Fractions of AluJ, AluS in AluY in enhancers and in RepeatMasker were compared using Fisher’s exact test (p-value < 2.2e-16 for the three Alu subfamilies). Importantly, these enhancers were overall depleted in Alu repeats, as previously reported by Andersson *et al.* [Nature, 2014]: We generated 63,285 random sequences using bedtools shuffle from F5.hg38.enhancers.bed and intersected the coordinates of the enhancers and these random sequences with those of the Alu repeats. We found that 5,089 enhancers and 11,967 random sequences intersected with at least one Alu repeat (Fisher’s exact test p-value < 2.2e-16, odds ratio = 2.7).

**Supplementary Figure S10.**
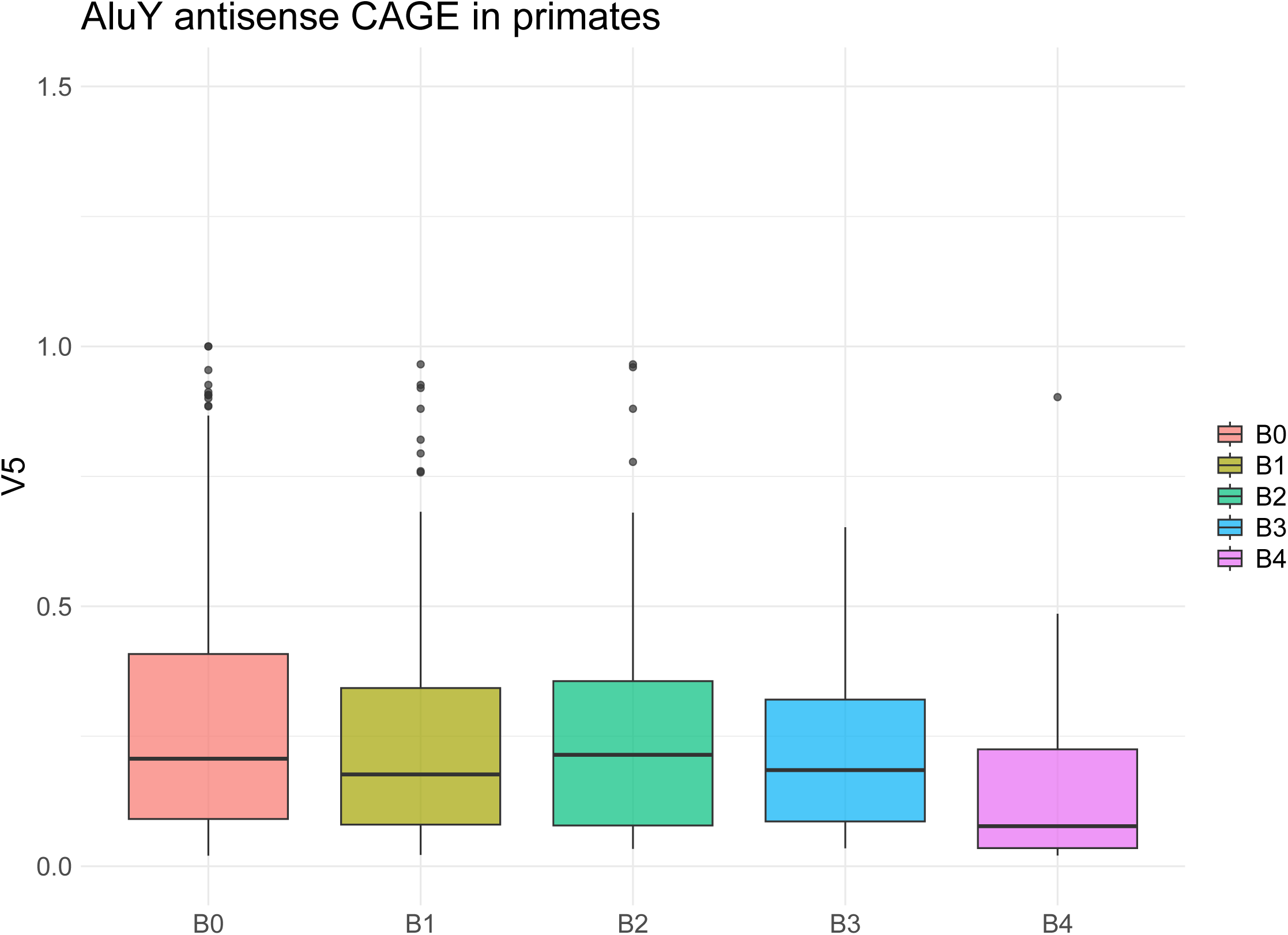
CAGE signal associated with STRs located in the antisense orientation of expressed primate-specific AluY elements as defined in [Zhang et al., 2019]. Number of transcribed STRs = 30 (for B4), 24 (for B3), 51 (for B2), 321 (for B1) and 363 (for B0). Wilcoxon test p-value = 3e-2 for CAGE signals associated with STRs located in the antisense orientation of Alu B4 and B3.

**Supplementary Figure S11.**
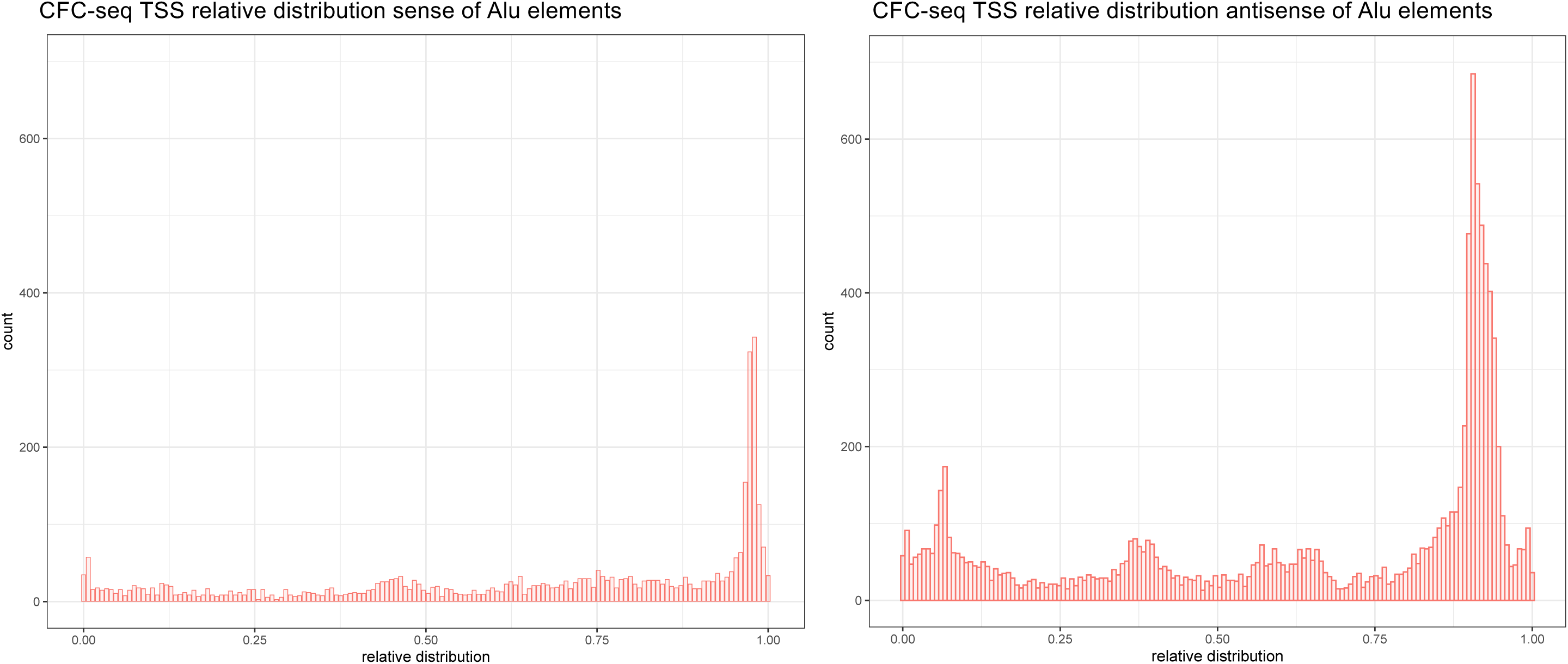
Relative distribution of CFC-seq TSSs in the sense (left) and antisense (right) orientation of Alu repeats as defined in RepeatMasker.

**Supplementary Figure S12.**
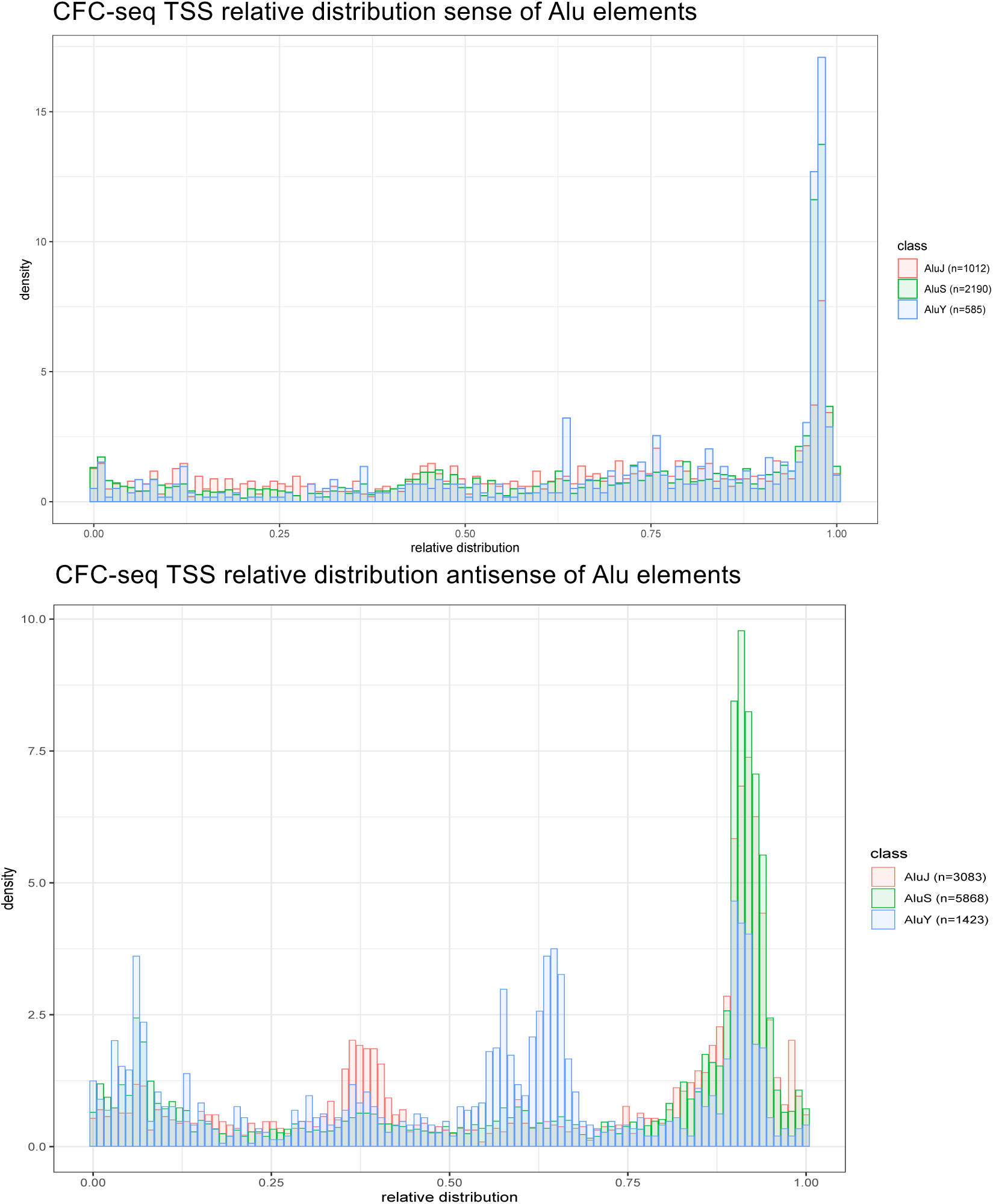
Relative distribution of transcript start sites (TSSs) in the sense (up) and antisense (bottom) orientation of AluJ (red), AluS (green) and AluY (blue) as defined by RepeatMasker. TSSs of annotated transcripts (ENCODE or FANTOM) and RNAs detected by CFC-seq are considered. The number of TSSs in each Alu class is indicated. Supplementary Figure S13 shows the same distribution considering only TSSs of RNAs detected during neuron differentiation of iPSC.

**Supplementary Figure S13.**
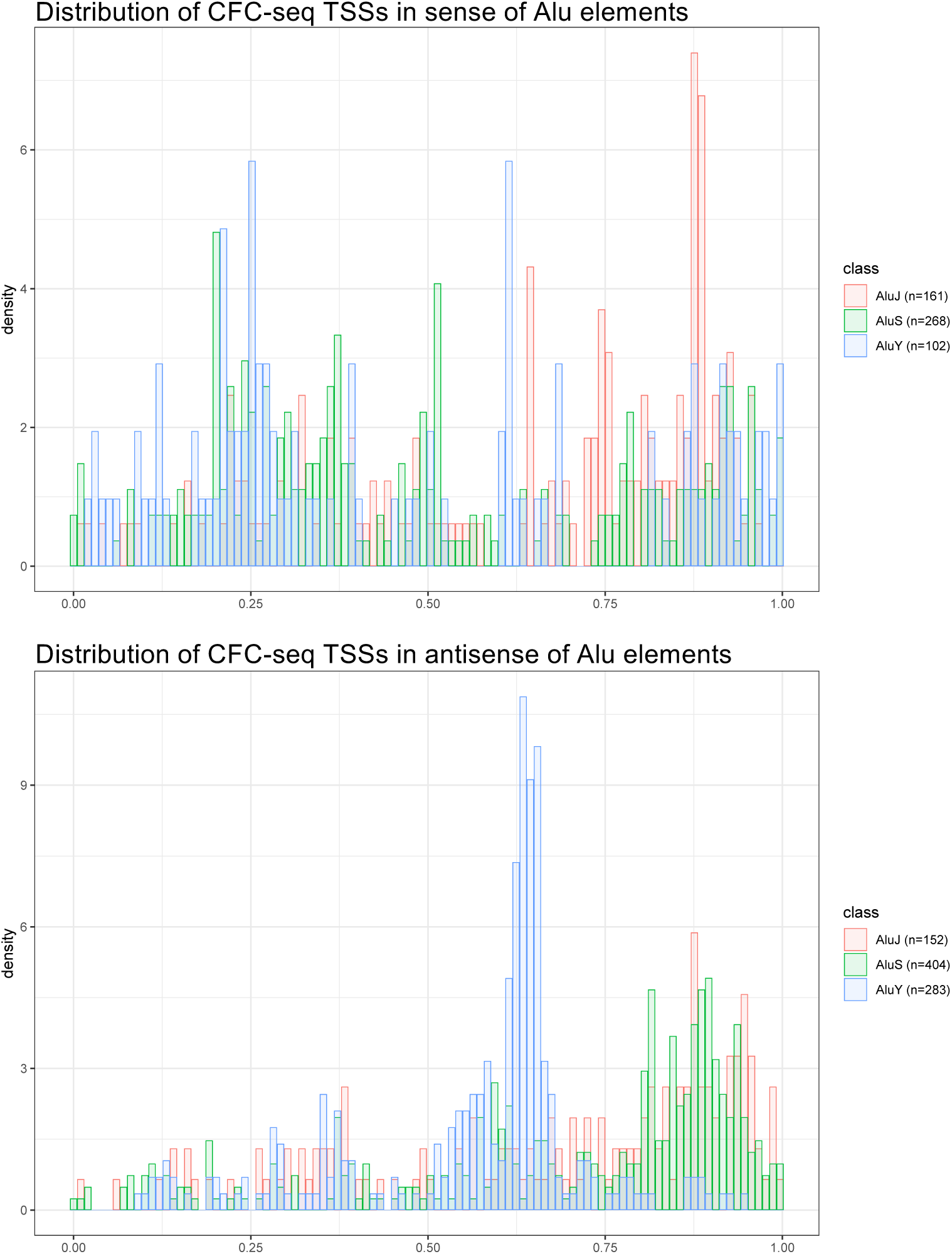
Relative distribution of CFC-seq TSSs in the sense (up) and antisense (bottom) orientation of AluJ (red), AluS (green) and AluY (blue) as defined by RepeatMasker. Only reads detected during the neuronal differentiation of iPSCs are considered. The number of TSSs in each Alu class is indicated.

**Supplementary Figure S14.**
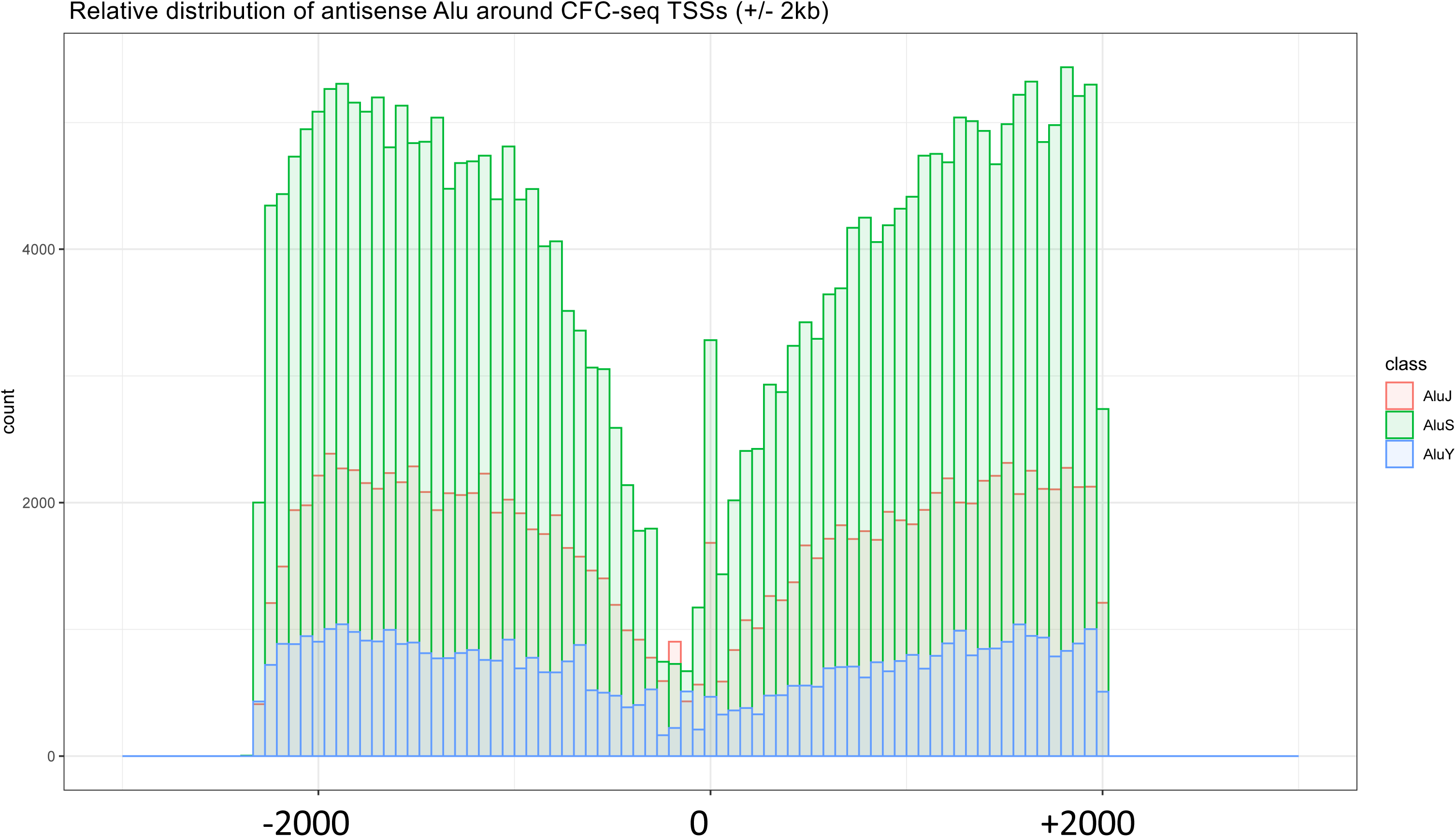
Distribution of 3’ends of RepeatMasker AluJ (red), AluS (green) and AluY (blue) around CFC-seq TSSs. (+/− 2kb). Only Alu located antisense of TSSs are shown. x-axis, distance to TSS

**Supplementary Figure S15.**
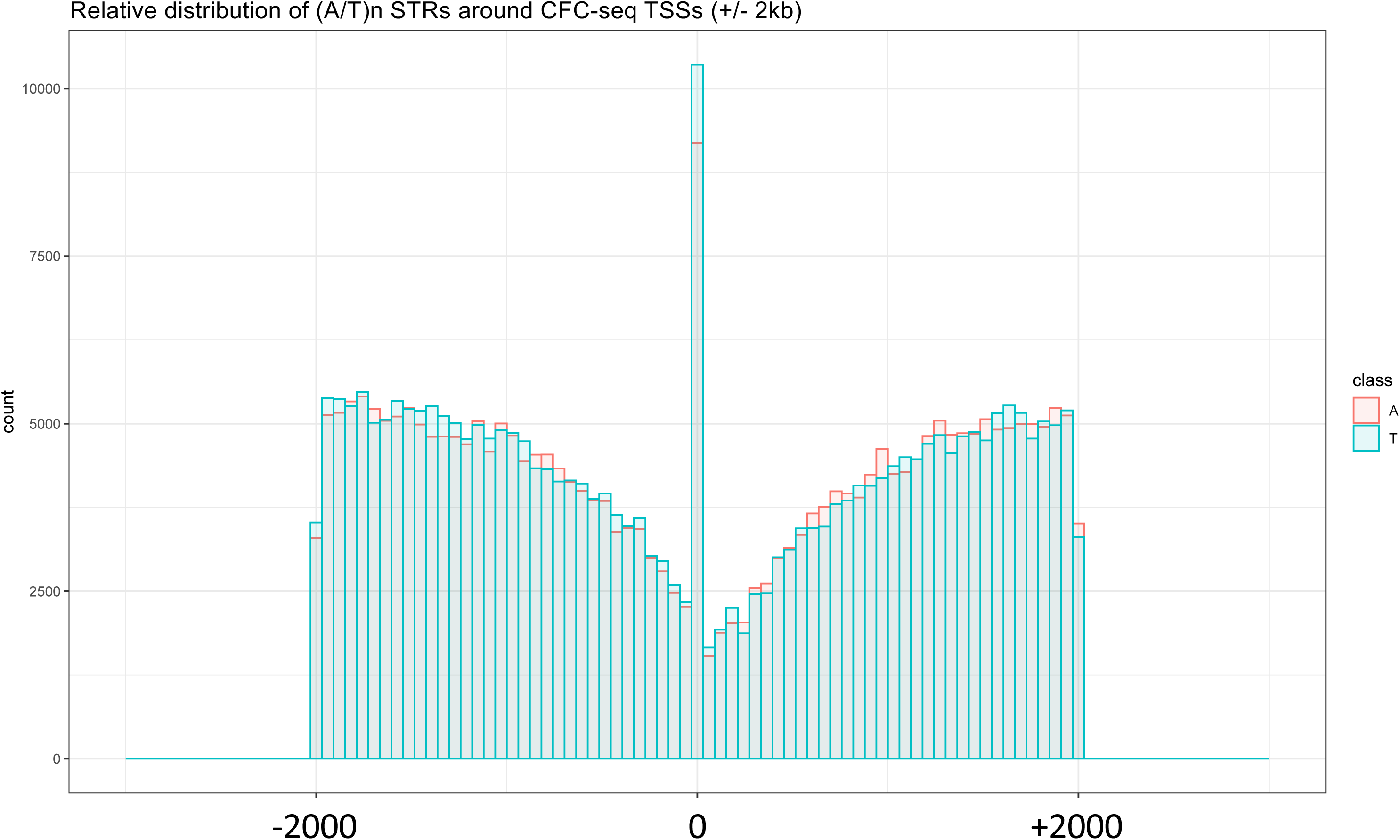
Distribution of (A)n (red) and (T)n STR (blue) midpositions around CFC-seq TSSs (+/−2kb). x-axis, distance to TS

**Supplementary Figure S16.**
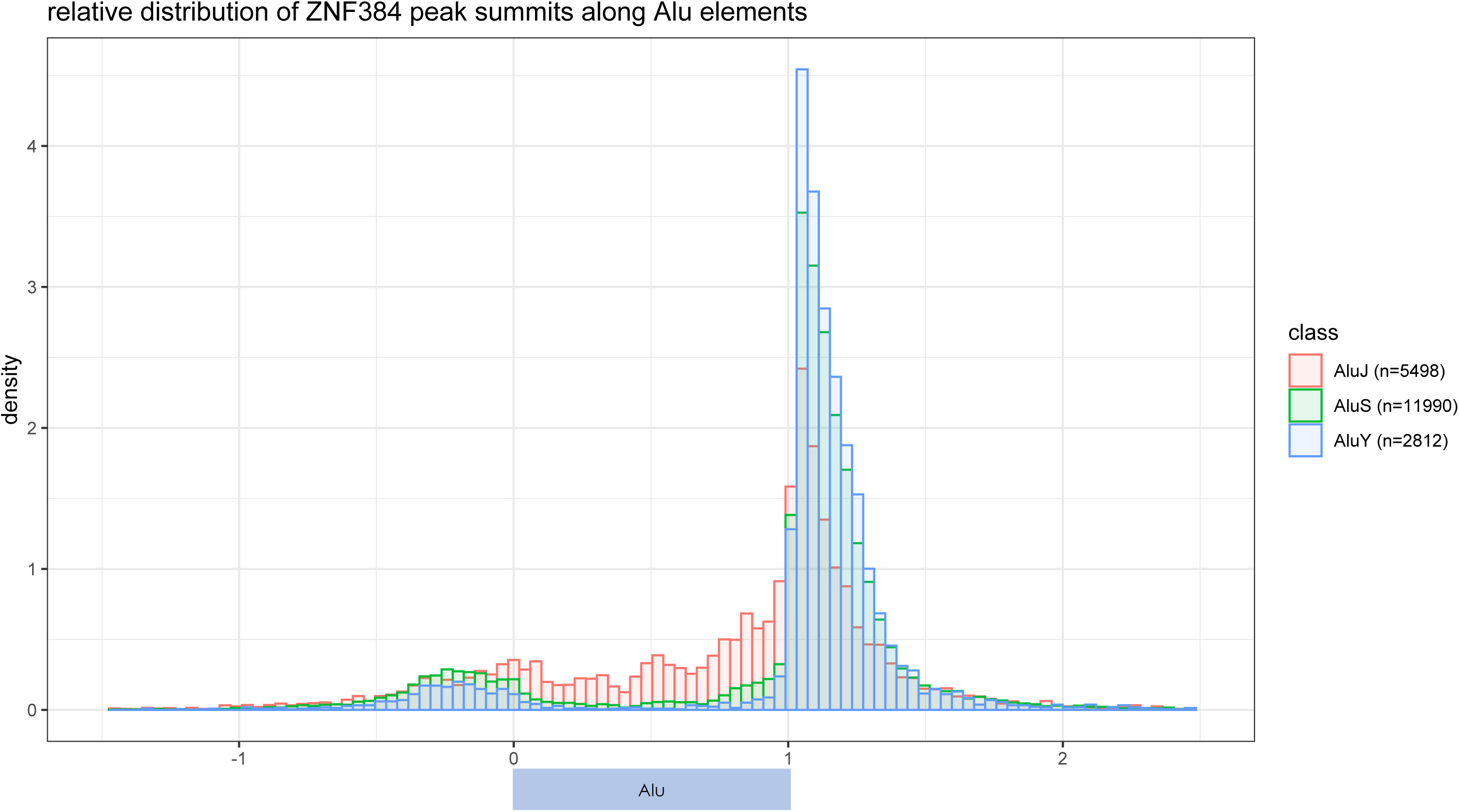
Relative distribution of ZNF384 ChIP peak summits along RepeatMasker AluJ (red), AluS (green) and AluY (blue). The number of ChIP peaks intersecting each Alu class is indicated.

